# Biomolecular Condensate Regulates Enzymatic Activity under Crowded Milieu: Synchronization of Liquid-Liquid Phase separation and Enzymatic Transformation

**DOI:** 10.1101/2022.06.16.496378

**Authors:** Bhawna Saini, Tushar Kanti Mukherjee

## Abstract

Cellular crowding plays a key role in regulating the enzymatic reactivity in physiological conditions, which is challenging to realize in the dilute phase. Enzymes drive a wide range of complex metabolic reactions with high efficiency and selectivity under extremely heterogeneous and crowded cellular environments. However, the molecular interpretation behind the enhanced enzymatic reactivity under a crowded milieu is poorly understood. Herein, using horseradish peroxidase (HRP) and glucose oxidase (GOx) cascade pair, we demonstrate for the first time that macromolecular crowding induces liquid-liquid phase separation (LLPS) via the formation of liquid-like condensates/droplets and thereby increases the intrinsic catalytic efficiencies of HRP and GOx. Both these enzymes undergo crowding induced homotypic LLPS via enthalpically driven multivalent electrostatic as well as hydrophobic interactions. Using a set of kinetic and microscopic experiments, we show that precise synchronization of spontaneous LLPS and enzymatic transformations is key to realize the enhanced enzymatic activity under the crowded environments. Our findings reveal an unprecedented enhancement (91–205-fold) in the catalytic efficiency (*k*_cat_/*K*_m_) of HRP at pH 4.0 within the droplet phase relative to that in the bulk aqueous phase in the presence of different crowders. In addition, we have shown that other enzymes also undergo spontaneous LLPS under macromolecular crowding, signifying the generality of this phenomenon under the crowded environments. More importantly, coalescence driven highly regulated GOx/HRP cascade reactions within the fused droplets have been demonstrated with enhanced activity and specificity under the crowded environments. The present discovery highlights the active role of membraneless condensates in regulating the enzymatic efficacy for complex metabolic reactions under the crowded cellular environments and may find significant importance in the field of biocatalysis.

## 1. Introduction

Cellular metabolic reactions are considerably more complex yet highly efficient than the in vitro reactions under dilute buffer conditions.^1^ How the complex and heterogeneous cellular environment regulates various complex biochemical reactions with high specificity and selectivity is a fundamental question in cell physiology that has not been understood yet. The cellular interior is highly crowded (~200–400 g/L) with various bio-macromolecules which occupy ~40% of its volume.^2^ Earlier, it has been demonstrated that macromolecular crowding plays a key role in regulating various fundamental processes including protein folding,^3–5^ aggregation,^6^ diffusion,^7,8^ thermal stability,^9^ DNA replication,^10^ and enzyme kinetics.^11–25^ In particular, the effect of crowding on enzymatic activity is complex and difficult to forecast. While some enzymes display enhanced activity^12,14,20^ in the presence of crowders, the catalytic rates of others are either decreased^11,13,15,17,22–25^ or remain unaltered.^26^ For example, Jiang and Guo observed more than 2-fold enhancement in the intrinsic catalytic efficiency of the isochorismate synthase in the presence of synthetic polymeric crowders as a consequence of significant reduction in the Michaelis-Menten constant (*K*_m_) via crowding induced conformational changes of the enzyme.^12^ Similarly, Ito and coworkers observed a significant jump in the enzymatic activity of endoplasmic reticulum glucosidase II in the presence of molecular crowders due to the crowding induced conformational change of the enzyme to its more active form.^20^ In another work, Norris and Malys reported a notable increase in the catalytic activity of glucose-6-phosphate dehydrogenase in the presence of different crowding agents, which has been explained on the basis of excluded volume theory.^14^ On the contrary, several studies have reported reduced activity of enzymes in crowded media due to the significant diffusion barriers of enzymes and substrates along with product inhibition in a highly crowded environment. ^11,13,15,17,22–25^ Moreover, crowding may also influence the magnitude of *K*_m_ by either lowering^12,13,15^ or increasing^8^ its value compared to that in dilute buffer conditions. In contrast, few earlier studies reported unaltered values of *K*_m_ upon crowding.^11,26^ The increase in *K*_m_ is accounted by considering diffusion resistance in a crowded environment, while the reduction is attributed to changes in activity coefficients of substrates and surrounding water molecules. These contrasting effects of macromolecular crowding on the kinetic parameters indicate the complex role of macromolecular crowding on the enzymatic reactions.

Despite these earlier efforts, the fundamental mechanism behind the activity enhancement upon crowding is still not clear. In general, the activity enhancement was accounted by considering stabilization of active conformation of enzymes via excluded volume effect under a crowded milieu. However, apart from the classical excluded volume effect, crowding can also modulate the soft protein-protein interactions via weak nonspecific chemical interactions.^27–32^ While the excluded volume effect is mainly entropic in nature and often stabilizes the compact native structure of proteins, nonspecific chemical interactions are enthalpic in nature and can either stabilize or destabilize the native protein structure. It is noteworthy to mention that most of the earlier studies on enzymatic kinetics in a crowded milieu have focused solely on the kinetic aspects while the influence of inert crowders on the physicochemical properties of enzymes is highly overlooked.

Another important aspect of macromolecular crowding is its ability to induce liquid-liquid phase separation (LLPS) of many disease-associated proteins containing intrinsically disordered regions (IDRs) with low complexity domains (LCDs) in their amino acid sequence via membraneless condensate/droplet formation.^33–37^ Very recently, we along with Maji and coworkers independently discovered that even globular proteins lacking any IDRs undergo LLPS upon crowding via soft protein-protein interactions.^38,39^ Although these recent findings delineate the role of soft protein-protein interactions on the physiological behavior of proteins, their impact on enzymatic activity is highly neglected in crowded environments. Inspired from our recent finding,^38^ we envisaged that multivalent soft intermolecular interactions between enzymes under a crowded milieu could potentially alter their secondary structures and subsequently modulate their metabolic reactions. Here we examined the catalytic activity of two well-known model enzymes, namely horseradish peroxidase (HRP) and glucose oxidase (GOx) in the absence and presence of inert crowders such as polyethylene glycol (PEG 8000), dextran 70, Ficoll 400, and bovine serum albumin (BSA) (Scheme 1), as their catalytic pathways were well explored in the literature.^40,41^

**Scheme 1.**
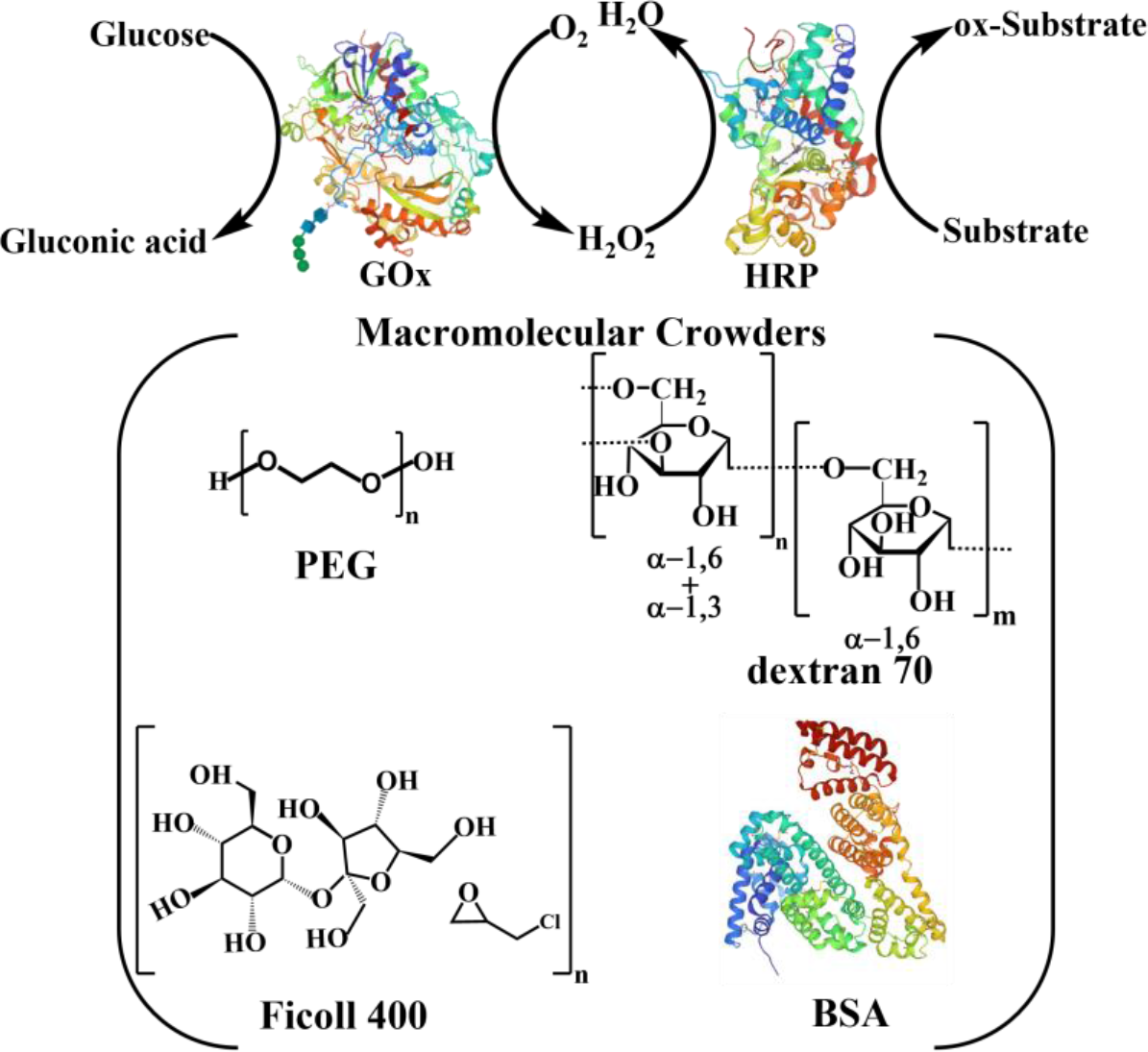
Schematics of Enzymatic Reactions and the Structures of Macromolecular Crowders.

More importantly, GOx/HRP cascade pair has found widespread applications in various fields including pharmaceuticals. ^42^ This pair catalyses the oxidation of glucose by GOx in the presence of oxygen (O_2_) to produce hydrogen peroxide (H_2_O_2_), which is subsequently utilized by HRP to oxidize organic or inorganic substrates by releasing O_2_ (Scheme 1). HRP belongs to the superfamily of heme-containing plant peroxidases with a molecular weight of 44 kDa and consists of a single polypeptide chain with 308 amino acid residues.^43^ On the other hand, GOx is a homodimer and belongs to the oxidoreductase family with a molecular weight of 160 kDa and consists of 605 amino acid residues.^44^ The primary objective of the present study is to thoroughly probe the physicochemical properties of HRP and GOx before, during and after the enzymatic transformations in the absence and presence of inert polymeric and protein crowders. Our present study revealed two fundamental discoveries which are highly relevant in the physiological context. First, we have discovered that both enzymes undergo spontaneous homotypic LLPS via the formation of liquid-like droplets under the crowded environment through multivalent electrostatic and hydrophobic interactions. Using two other control enzymes, we have illustrated that LLPS may be a common phenomenon for these classes of biocatalysts in a crowded environment. Second, we have discovered that precise synchronization of LLPS and enzymatic transformation is critical to boost the intrinsic catalytic activity of phase separated enzymes inside the droplet phase. These discoveries are physiologically relevant in the sense that nature may also utilizes similar spatio-temporal synchronization of these complex events for optimum activity and selectivity under highly heterogeneous and crowded subcellular compartments.

## 2. Results and Discussion

### 2.1. Macromolecular Crowding Induces LLPS of HRP and GOx

Recent studies have highlighted that inert macromolecular crowders can effectively modulate the soft protein-protein interactions of ordered proteins and induce LLPS via the formation of liquid-like droplets.^33–39^ Inspired from these findings, we initially seek to know whether the presence of inert crowders could alter the intermolecular protein-protein interactions of two well-known model enzymes, namely HRP and GOx under physiological conditions. In the present study, protein-protein interactions were studied by using confocal laser scanning microscopy (CLSM) and field-emission scanning electron microscopy (FESEM). CLSM measurements were performed using 0.5 μM fluorescently labeled enzymes (see SI for more details). HRP and GOx were labeled with rhodamine B isothiocyanate (RBITC) and fluorescein isothiocyanate (FITC) to visually monitor them under the confocal microscope. Notably, the aqueous solutions of 0.5 μM HRP and GOx in the absence and presence of crowders remain isotropic in nature and no visible turbidity appears within 2 days of incubation at 37 °C (Figure S1). However, the aqueous solution of GOx in the presence of 10% PEG shows visible turbidity after 3 days of incubation at 37 °C. Surprisingly, both enzymes in the presence of 10% PEG 8000 at 37 °C display uniform spherical assemblies as revealed from confocal differential interference contrast (DIC) images captured after 1 h of incubation (Figure 1A). No such assemblies are observed in the absence of PEG (Figure 1B), indicating the active role of crowders behind the formation of these spherical assemblies. CLSM images with RBITC-labeled HRP and FITC-labeled GOx confirm the presence of enzymes inside these assemblies (Figure 1C). It is noteworthy to mention that fluorescent labeling does not perturbs the formation and morphologies of these assemblies as revealed from our control experiments with 10% and 100% labeled enzymes (Figure S2). These assemblies exhibit liquid-like droplet characteristics such as fusion of smaller droplets into a bigger droplet (spontaneous merging upon contact), dripping (separation of smaller droplet from a larger droplet), and surface wetting (spreading on the surface) (Figure 1C). Similar liquid-like droplet formation via LLPS was observed previously for a wide range of biomolecules.^33–39^ To know whether this phenomenon is due to any specific interactions with PEG, we used other inert crowders to mimic the macromolecular crowding effect of PEG.

**Figure 1.**
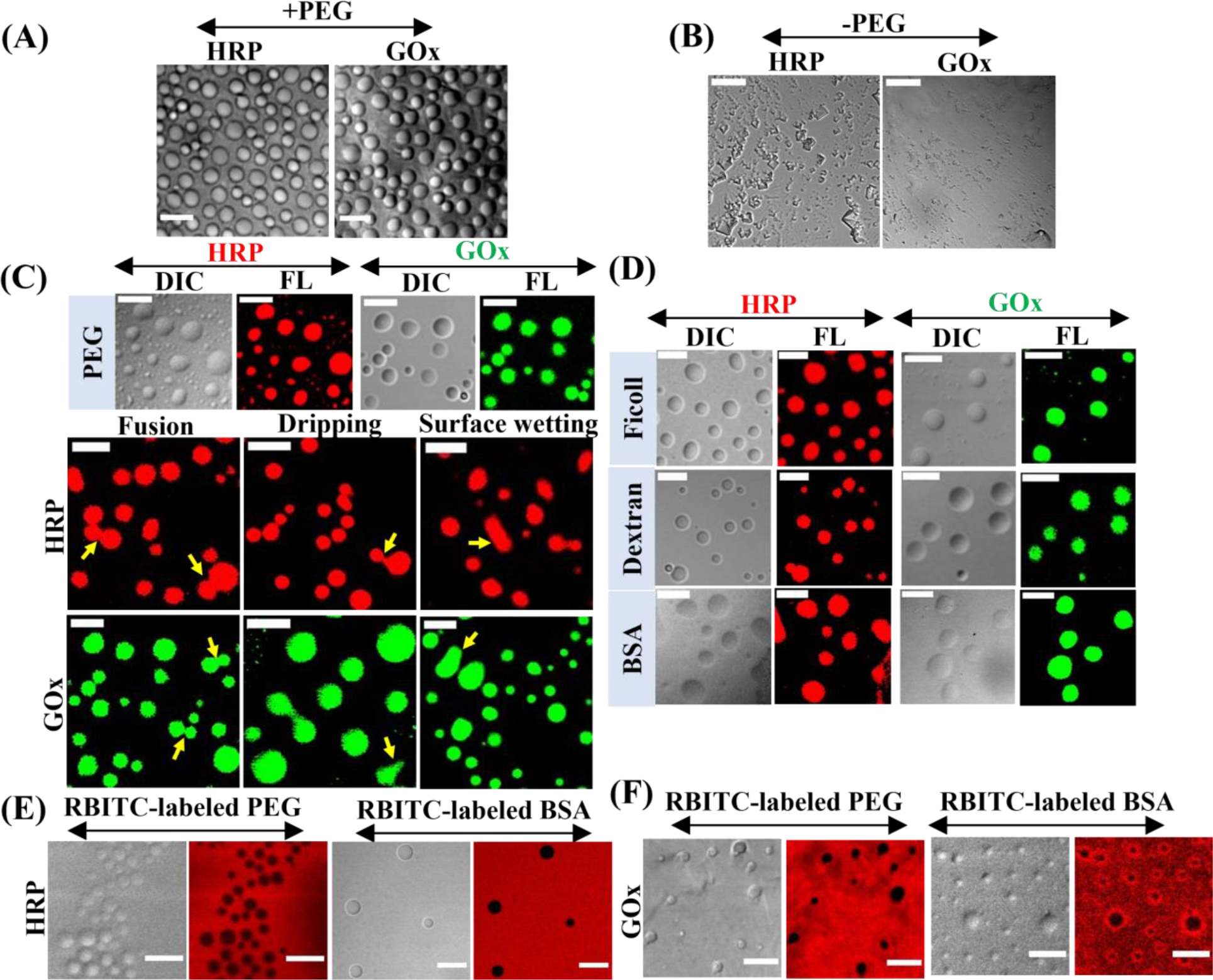
DIC images of HRP and GOx in the (A) presence, and (B) absence of 10% PEG. (C) Confocal images of RBITC-labeled HRP and FITC-labeled GOx droplets formed in the presence of 10% PEG showing coalescence, dripping, and surface wetting phenomena. (D) Confocal images showing droplets of HRP and GOx in the presence of 12.5% Ficoll, 10% dextran, and 20 mg/mL BSA. Confocal images of (E) HRP, and (F) GOx droplets in the presence of RBITC-labeled mPEG-NH_2_ and RBITC-labeled BSA. The scale bars correspond to 5 μm.

Notably, droplet formation has also been observed in the presence of 12.5% Ficoll 400, 10% dextran 70, and 20 mg/mL BSA, suggesting a general crowding effect (Figure 1D). Independent experiments with FESEM measurement also confirm the formation of droplets in the presence of polymeric crowders (Figure S3). These droplets are colloidally stable over a period of 15 days without any structural distortion (Figure S4). To know whether these droplets are formed via homotypic (only enzymes are involved) or heterotypic (both enzyme and crowder are involved) LLPS phenomenon, we utilized fluorescently labeled PEG (RBITC-labeled mPEG-NH_2_, MW 5000) and BSA (RBITC-labeled) to visualize their partition using CLSM. These sets of experiments were performed with unlabeled enzymes. It is evident that both PEG and BSA are excluded from the phase-separated droplets of HRP and GOx as revealed from their background fluorescent signals (Figure 1E,F). These observations authenticate that the droplets of HRP and GOx are formed via homotypic LLPS, where crowders are totally excluded from the droplet phase. Here it is important to note that the present homotypic LLPS phenomenon is fundamentally different than the aqueous two-phase system (ATPS) reported earlier with PEG/sodium citrate and PEG/dextran aqueous mixtures.^22,25^ To show that PEG alone in an aqueous buffer does not yield any ATPS, we added free FITC dye in the aqueous mixture for visual detection. The daylight photograph clearly reveals the isotropic mixture of PEG and FITC in the aqueous buffer (Figure S5). However, addition of 220 mM sodium citrate indeed results in ATPS (Figure S6), as reported earlier. Taken together, these findings reveal that both these enzymes undergo spontaneous homotypic LLPS in the presence of inert crowders via the formation of liquid-like droplets.

Next, we seek to address the physicochemical properties of these phase-separated droplets as a function of incubation time at 37 °C. CLSM images were captured for samples with varying incubation times of 5, 15, 30, and 60 min. Figure 2A shows the confocal images of HRP and GOx droplets at different incubation times.

**Figure 2.**
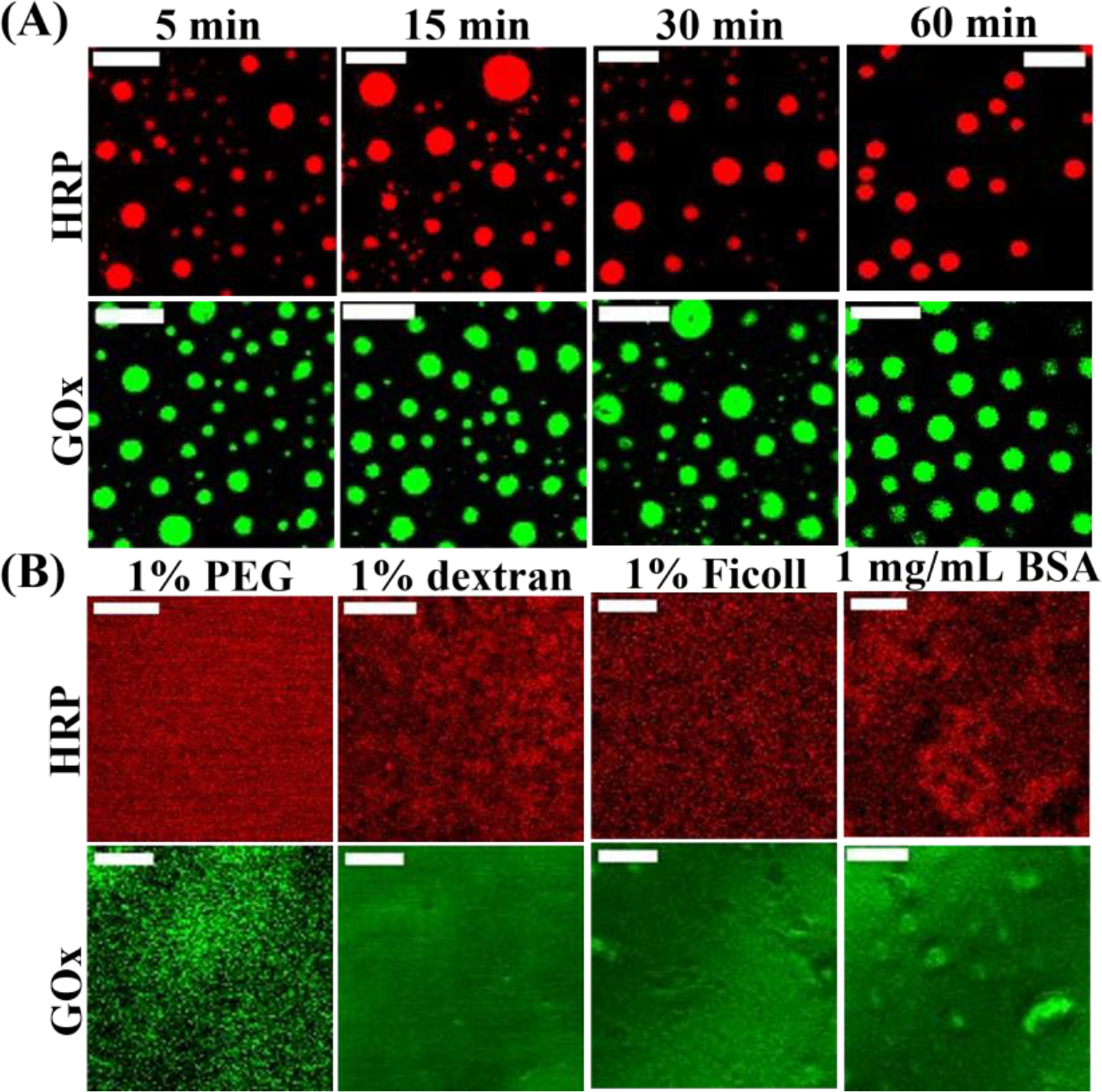
(A) Confocal images of RBITC-labeled HRP and FITC-labeled GOx droplets in the presence of 10% PEG as a function of incubation time. (B) Confocal images showing absence of any droplets in the presence of 1% PEG, 1% dextran, 1% Ficoll, and 1 mg/mL BSA. The scale bars correspond to 5 μm.

It is evident that irrespective of the incubation time well-dispersed droplets have been observed for both the enzymes. However, droplets become homogeneous in size upon an increase in the incubation time due to spontaneous fusion events and after 60 min of incubation, droplets of both the enzymes exhibit uniform size distribution (Figure 2A). Importantly, droplet formation is inhibited completely at lower concentrations of crowders, indicating the critical role of crowders to induce LLPS (Figure 2B).

This intriguing observation of crowding induced LLPS of these two model enzymes is quite surprising to us initially as this phenomenon was not anticipated in earlier studies although the effect of crowding on the kinetics of both these enzymes was extensively investigated. In general, proteins having IDRs with LCDs in their amino acid sequence exhibit spontaneous LLPS beyond a critical concentration in the absence or presence of macromolecular crowders.^33–39,47–49^ However, recent findings revealed that the presence of IDRs with LCDs is not an essential requirement for biomacromolecules to undergo LLPS.^38,39^ To know whether these two model enzymes contain any LCDs and IDRs, we used sequence prediction algorithms, namely Simple Modular Architecture Research Tool (SMART)^45^ and IUPred2,^46^ respectively. While SMART analysis of GOx sequence reveals three LCDs (residues 4–17, 47–61, and 363–371), IUPred2 disorder prediction algorithm predicts around six short disorder regions (residues 127–130, 192–204, 222–226, 237–243, 287–291, and 359–360) (Figure S7). However, neither LCDs nor any disorder regions are predicted for HRP (Figure S8), suggesting that the presence of LCDs and/or IDRs is not an essential prerequisite for LLPS. Therefore, our observations suggest that macromolecular crowders induce homotypic LLPS of both these enzymes via intermolecular interactions between short patches of polypeptide chains. Next, we explored the molecular origin of these intermolecular protein-protein interactions under the crowded environment.

### 2.2. Nature of Intermolecular Interactions

#### Effect of temperature and pH

LLPS of biomolecules is mainly driven by various weak multivalent intermolecular interactions including electrostatic, hydrophobic, hydrogen bonding, aromatic and/or cation-π interactions.^33–39,47–49^ To decipher the precise role of these interactions, we studied the droplet formation pathways of HRP and GOx as a function of temperature, pH, and ionic strength under CLSM and FESEM in the presence of 10% PEG. Previous studies have highlighted the critical role of temperature on the feasibility of LLPS of a wide range of proteins.^33,35,38,39,49^ Depending on the nature of intermolecular interactions, LLPS of biomacromolecules exhibits either upper critical solution temperature (UCST)^38^ or lower critical solution temperature (LCST)^35^ or both^49^. The feasibility of LLPS of both the enzymes in the presence of 10% PEG was monitored by varying the temperatures in the range of 4–90 °C using CLSM and FESEM measurements (Figure 3A and Figure S9). It has been observed that droplets of both the enzymes are stable in the temperature range of 4–80 °C; however, droplet formation is completely inhibited at 90 °C. These observations suggest that LLPS of both the enzymes follow UCST profile. Similar UCST profile has been reported earlier for other proteins.^38,49^ The observed UCST profiles indicate a dominant role of enthalpically driven intermolecular interactions over the entropy of mixing as described previously by the Flory– Huggins theory.^50,51^

**Figure 3.**
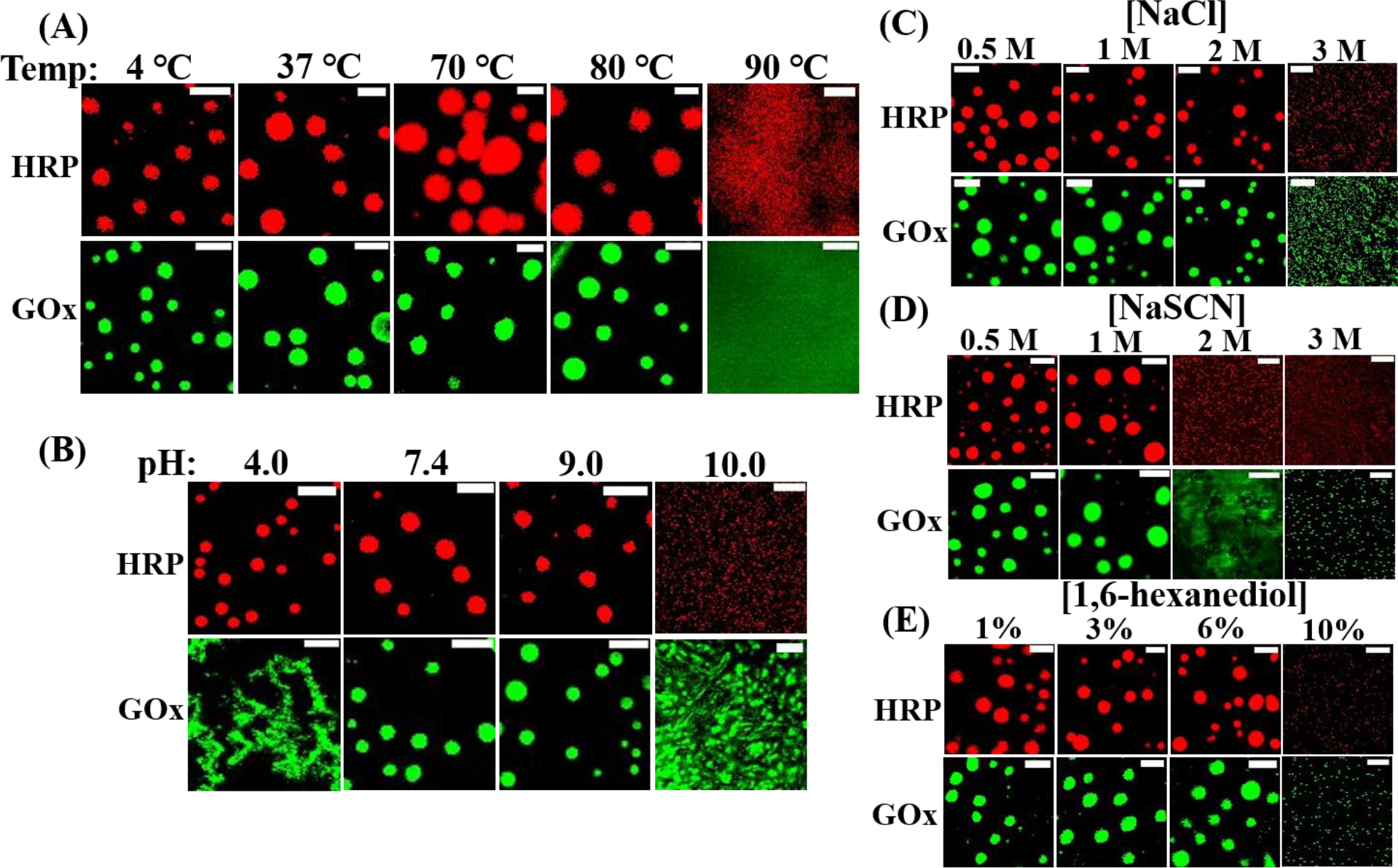
Confocal images showing the stability of RBITC-labeled HRP and FITC-labeled GOx droplets as a function of (A) temperature, (B) pH, (C) NaCl concentrations, (D) NaSCN concentrations, and (E) 1,6-hexanediol concentrations. The scale bars correspond to 5 μm.

The pH is also known to influence the LLPS profiles of proteins via modulation of charges and/or secondary structures. Next, the feasibility of LLPS of both the enzymes in the presence of 10% PEG is investigated as a function of solution pH in the range of 4–10 (Figure 3B and Figure S10). HRP in the presence of 10% PEG forms well-dispersed droplets in the pH range of 4–9; however, droplet formation is inhibited at pH 10 as revealed from CLSM images. In contrast, droplets of GOx are stable in the pH range of 7.4–9; however, they disintegrate at pH values of 4 and 10 (Figure 3B and Figure S10). These findings indicate the active role of charged residues of HRP and GOx behind the droplet formation mechanism by considering their pI values of 8.8 and 4.2, respectively.^52^

#### Effect of salt and aliphatic alcohol

Intermolecular protein-protein interactions can be modulated to a great extent by varying the concentrations of salts and aliphatic alcohols. Next, we tested the feasibility of LLPS in the presence of 10% PEG upon the addition of varying concentrations of a neutral salt, sodium chloride (NaCl), chaotropic salt, sodium thiocyanate (NaSCN), and aliphatic alcohol, 1,6-hexanediol. Droplets of both the enzymes are found to be stable up to a NaCl concentration of 2M (Figure 3C and Figure S11). However, droplet formation is completely inhibited in the presence of 3 M NaCl for both the enzymes, indicating the key role of electrostatic intermolecular interactions behind the LLPS of both the enzymes.^33–35,38,48,49^ It should be noted that LLPS solely driven by electrostatic interactions can be inhibited at much lower NaCl concentration (<500 mM).^33,48,49^ This suggests that the present LLPS of HRP and GOx in the presence of 10% PEG may not be solely due to the electrostatic intermolecular interactions. To know whether any hydrophobic protein-protein interactions play any role in the observed LLPS of HRP and GOx, we varied the concentrations of NaSCN and 1,6-hexanediol, which are known to disrupt hydrophobic protein-protein interactions.^34,38,48^ Droplets of both enzymes remain intact up to 1M of NaSCN; however, droplet formation is inhibited completely at and beyond 2 M of NaSCN (Figure 3D and Figure S12). Similarly, droplets of both enzymes remain intact in the presence of 1-6% of 1,6-hexanediol; however, they disintegrate completely at 10% of 1,6-hexanediol (Figure 3E and Figure S13). These findings authenticate the active role of hydrophobic protein-protein interactions behind the observed LLPS of HRP and GOx in the presence of crowders. Taken together, our findings illustrate that the droplet formation of both enzymes via LLPS is primarily driven by multivalent electrostatic as well as hydrophobic intermolecular interactions between short patches of polypeptide chains. Moreover, the preset findings reveal that macromolecular crowding effectively favors these soft protein-protein interactions and drives the LLPS under the physiological conditions.

### 2.3. Alteration of Secondary Structures of Enzymes

To know the conformational perturbation of HRP and GOx in the presence of crowders, we performed circular dichroism (CD) measurements in the absence and presence of 10% PEG. The far-UV CD spectrum of HRP at pH4.0 displays two minima at 208 and 222 nm, suggesting an α-helical rich secondary structure(Figure 4A).

**Figure 4.**
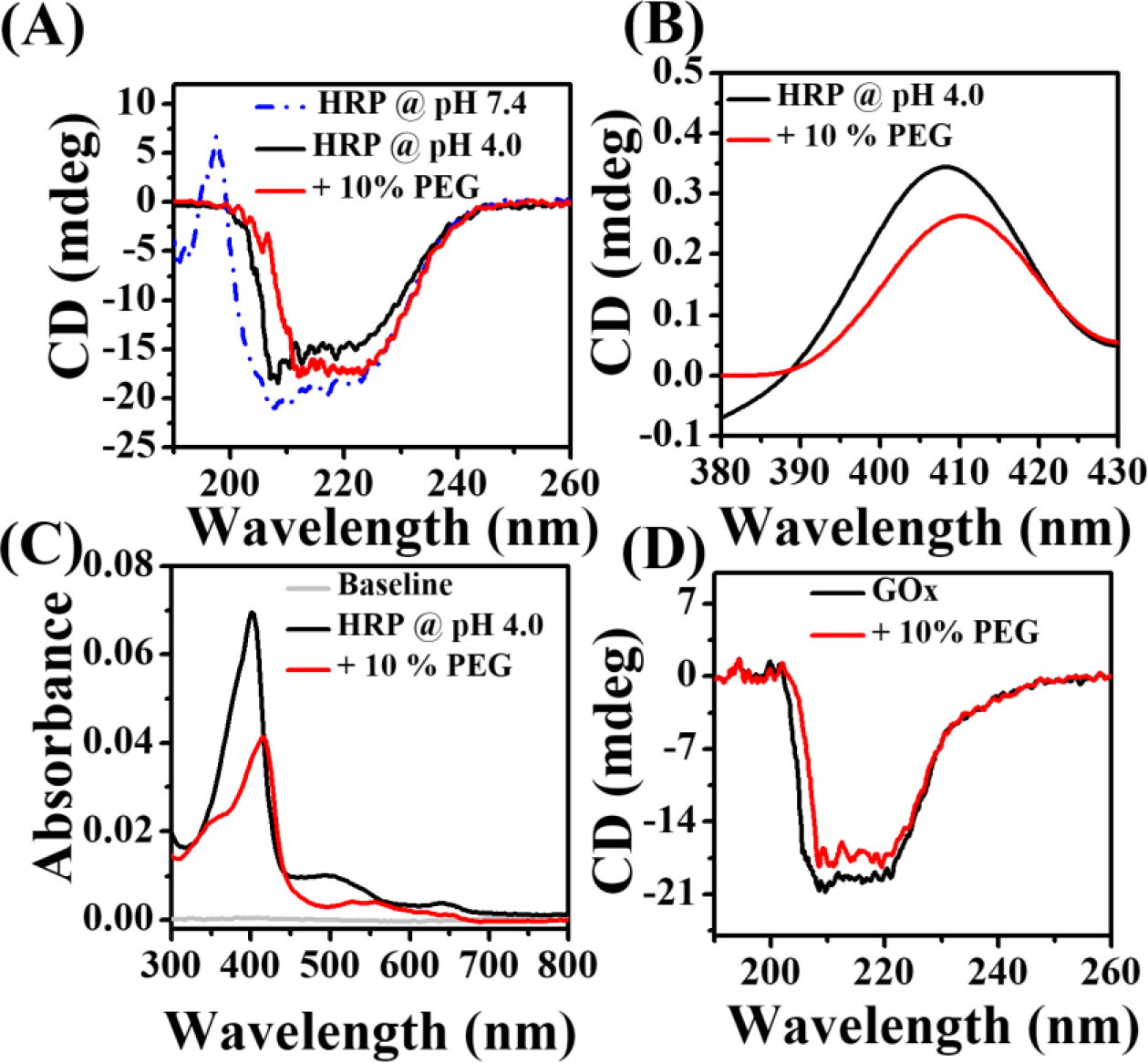
Effect of 10% PEG on (A) far-UV CD, (B) Soret region CD, and (C) UV-vis absorption spectra of 0.5 μM HRP. (D) Changes in the far-UV CD spectra of 0.5 μM GOx in the presence of 10% PEG.

The spectrum of HRP at pH4 differs slightly from that at pH 7.4 with decrease in ellipticity at 222 nm (Figure 4A, dotted line), indicating pH-dependent conformation change. Importantly, a distinct red shift of 4 nm with an increase in ellipticity at 222 nm has been observed in the presence of PEG. Similarly, the CD spectrum at the Soret region and UV-vis spectrum of HRP in the presence of PEG display a prominent red shift of 3 and 15 nm, respectively compared to that in the aqueous buffer (Figure 4B, C). These spectral change of HRP upon crowding indicates more compact secondary structure and higher integrity of the heme pocket.^53^ In contrast, the ellipticity of GOx at 222 nm decreases slightly upon crowding (Figure 4D), suggesting a subtle conformational change of GOx. Therefore, these findings clearly authenticate the altered secondary structures of phase separated enzymes under the crowded environment relative to that in the aqueous buffer. Next, we seek to address how these conformationally altered phase separated enzymes in a crowded milieu drive their respective catalytic transformations. Here it is important to mention that the present study is first of its kind to demonstrate the detailed kinetic aspects of enzymatic transformations considering the phase separation in a crowded milieu, which is completely ignored in earlier reports.

### 2.4. Effect of LLPS on the Enzymatic Kinetics

For all the kinetic experiments, we kept the enzyme concentration fixed at 25 pM as at high concentration (0.5 μM), the enzymatic transformations were too fast to monitor in the crowded environments. Control experiments with 25 pM of enzymes reveal similar kinds of LLPS in the presence of 10% PEG (Figure S14). The kinetics of HRP and GOx catalyzed reactions were monitored using a UV-vis spectrophotometer at their optimum pH values of 4 and 7.4, respectively (Figure S15). Different sets of kinetic experiments were performed under distinct conditions to highlight the overall effect of crowding on the enzymatic activity. We mainly monitored the enzymatic transformations before and after the phase separation of HRP and GOx in the presence of different crowders such as 10% PEG 8000, 10% dextran 70, 12.5% Ficoll 400, and 20 mg/mL BSA. Moreover, the enzymatic kinetics of HRP were monitored after the LLPS in a time-dependent manner to highlight the effect of diffusion and substrate partitioning inside the phase separated droplets in a crowded milieu. Finally, the feasibility of GOx/HRP cascade reaction was investigated in the presence of 10% PEG and 20 mg/mL BSA after their phase separation using different substrates such as 3,3́ʹ,5,5ʹ-tetramethylbenzidine (TMB), o-phenylenediamine (OPD), and 2,2ʹ-azino-bis(3-ethylbenzothiazoline-6-sulfonic acid (ABTS).

#### Enzymatic Kinetics before Phase Separation

Initially, we tested the influence of various crowders on the individual enzymatic kinetics of HRP and GOx just after their addition in pH 4.0 and 7.4 aqueous buffers, respectively (Figure 5A).

**Figure 5.**
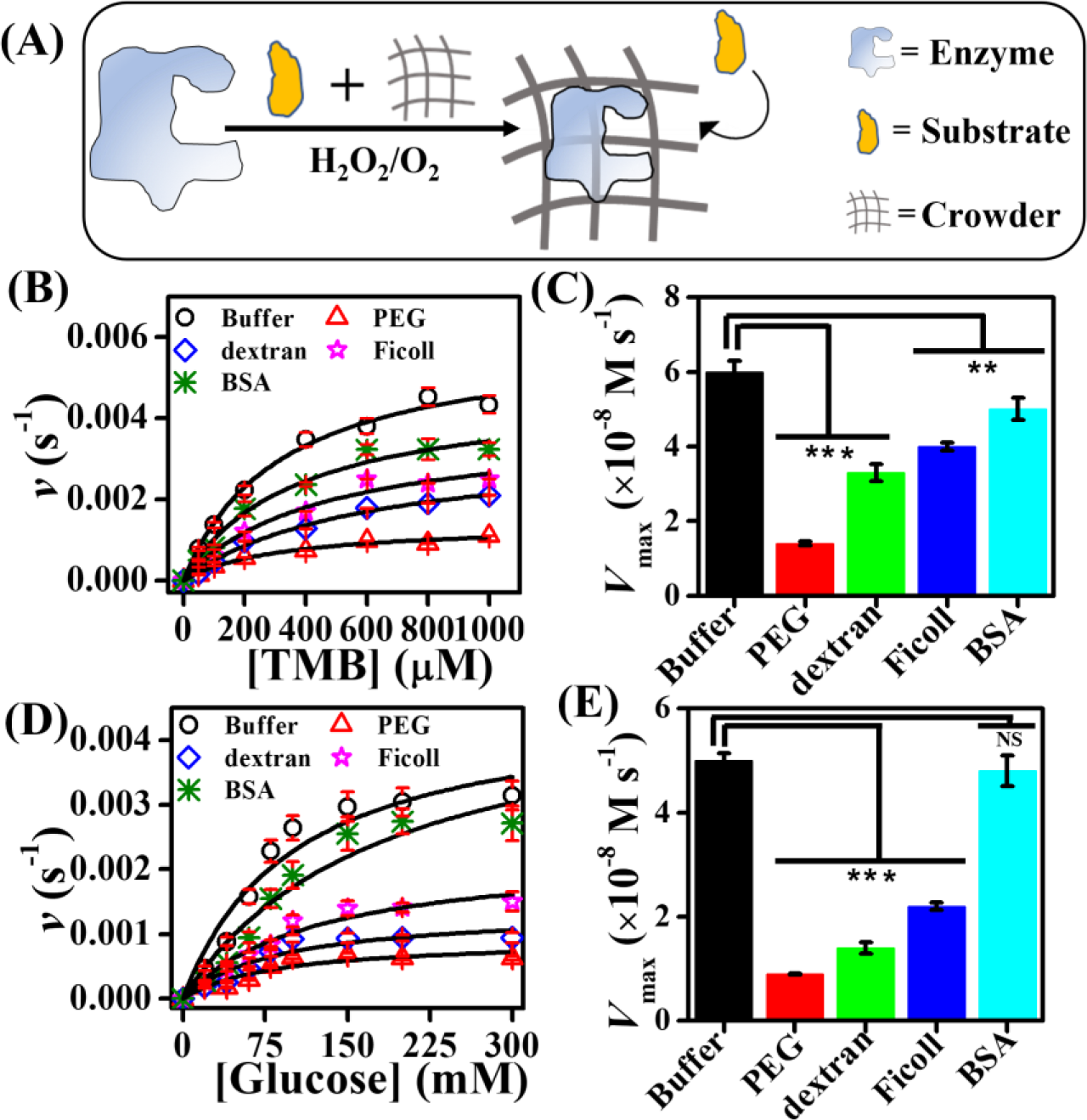
(A) Illustration of enzymatic reaction in the presence of substrates and crowder. The effect of different crowders on the (B) Michaelis-Menten plots of HRP, (C) *V*_max_ values of HRP,(D) Michaelis-Menten plots of GOx, and (E) *V*_max_ values of GOx. The data represent the mean ± s.e.m. for three (n = 3) independent experiments. Statistical significance was assessed by a two-tailed, unpaired Student’s *t*-test with^* * *^ *P* value <0.001, ^* *^ *P* value <0.01 and not significant (NS; *P*>0.05).

HRP catalyzed reactions were monitored using TMB as a substrate by recording the changes in the absorbance at 650 nm as a function of reaction time (Figure S16). The enzymatic rate of 25 pM HRP in the presence of 8.8 mM H_2_O_2_ in the aqueous acetate buffer (pH 4.0) at 37 °C follows a typical Michaelis-Menten plot as a function of TMB concentrations (Figure 5B). Notably, the rate decreases drastically in the presence of different crowders as revealed from the Michaelis-Menten plots (Figure 5B). The observed rate is least in 10% PEG followed by 10% dextran, 12.5% Ficoll, and 20 mg/mL BSA. These data were fitted with the Michaelis-Menten equation to obtain maximum velocity (*V*_max_) and Michaelis-Menten constant (*K*_m_). The calculated *V*_max_ values for different crowders are compared in Figure 5C. It is evident that *V*_max_ decreases in the presence of crowders and more so in the presence of polymeric crowders. Previously, it has been shown that macromolecular crowding slowdown enzymatic kinetics significantly.^11,13,15,17,22–25^ For example, Keating and coworkers demonstrated lower activity of HRP at pH 7.4 in the presence of PEG and dextran due to the crowding induced conformational change and lowered diffusion coefficients of substrates in viscous polymeric solution.^21^ While they observed an increase in the *K*_m_ value, the *V*_max_ value of HRP decreased in the presence of crowders. Moreover, it is known that polymeric crowders increase the viscosity of the solution more compared to the protein crowders and as a result the rate of the diffusion-controlled reactions slowdown significantly in the presence of polymeric crowders.^24^ The estimated *K*_m_ and turnover numbers (*k*_cat_) are tabulated in Table 1. While the *K*_m_ of TMB remains unaltered in the presence of PEG, a noticeable increase has been observed in the presence of dextran, Ficoll, and BSA. The estimated *K*_m_ values are in good agreement with the previous report.^54^ On the other hand, *k*_cat_ decreases from a value of 2.4 × 10^3^ in buffer to 5.6 × 10^2^, 1.3 × 10^3^, 1.6 × 10^3^, and 2.0 × 10^3^ s^−1^ in the presence of 10% PEG, 10% dextran, 12.5% Ficoll, and 20 mg/mL BSA, respectively.

**Table 1.** Michaelis-Menten Parameters Estimated Instantly in Different Solutions.

| | $K_m$ (mM) | | $k_{cat}$ (s<sup>-1</sup>) | |
| --- | --- | --- | --- | --- |
|  | HRP | GOx | HRP | GOx |
| Buffer | $0.30 \pm 0.06$ | $101.0 \pm 5.2$ | $(2.4 \pm 0.07) \times 10^3$ | $(2.0 \pm 0.01) \times 10^3$ |
| 10% PEG | $0.30 \pm 0.05$ | $83.3 \pm 2.3$ | $(5.6 \pm 0.5) \times 10^2$ | $(3.7 \pm 0.1) \times 10^2$ |
| 10% dextran | $0.60 \pm 0.05$ | $88.0 \pm 1.2$ | $(1.3 \pm 0.07) \times 10^3$ | $(5.6 \pm 0.8) \times 10^2$ |
| 12.5% Ficoll | $0.40 \pm 0.03$ | $84.4 \pm 1.6$ | $(1.6 \pm 0.1) \times 10^3$ | $(8.8 \pm 0.7) \times 10^2$ |
| 20 mg/mL BSA | $0.40 \pm 0.07$ | $84.0 \pm 2.1$ | $(2.0 \pm 0.04) \times 10^3$ | $(1.9 \pm 0.1) \times 10^3$ |

Similar trends have been observed for GOx kinetics with glucose as a substrate in the presence of different crowders (Figure 5D and Figure S17). The *V*_max_ decreases significantly in the presence of polymeric crowders compared to that in the aqueous buffer (Figure 5E). Moreover, a noticeable decrease in the *K*_m_ value of glucose has been observed in the presence of crowders (Table 1). The estimated *K*_m_ values are in good agreement with the previous report.^55^ The lower values of *K*_m_ in the presence of crowders indicate higher binding affinity of glucose possibly due to the altered conformation of GOx in the crowded environments. While the estimated *k*_cat_ of GOx decreases remarkably in the presence of polymeric crowders, it remains almost unaltered in the presence of 20 mg/mL BSA. Taken together, our findings reveal that the enzymatic kinetics of HRP and GOx slowdown appreciably in the crowded environment, similar to the previous reports. ^11,13,15,17,22–25^ Notably, these observations are valid only when the enzymatic reactions are monitored instantly after the addition of enzymes in the crowded environment. However, these enzymes can simultaneously undergo LLPS during the course of these transformations and hence the present framework is not an ideal platform to explore the effect of macromolecular crowding. To know the overall impact of crowding on the enzymatic kinetics, we next bifurcated the complete process into two independent events, namely LLPS followed by enzymatic transformations.

#### Enzymatic Kinetics After Phase Separation

Here, we first allowed the enzymes to undergo LLPS in the presence of crowders. Enzymes were incubated in the aqueous buffers in the presence of crowders at 37 °C for 1 h to have a homogeneous growth of phase separated droplets. Subsequently, enzymatic reactions were monitored instantly after the addition of substrates (Figure 6A).

**Figure 6.**
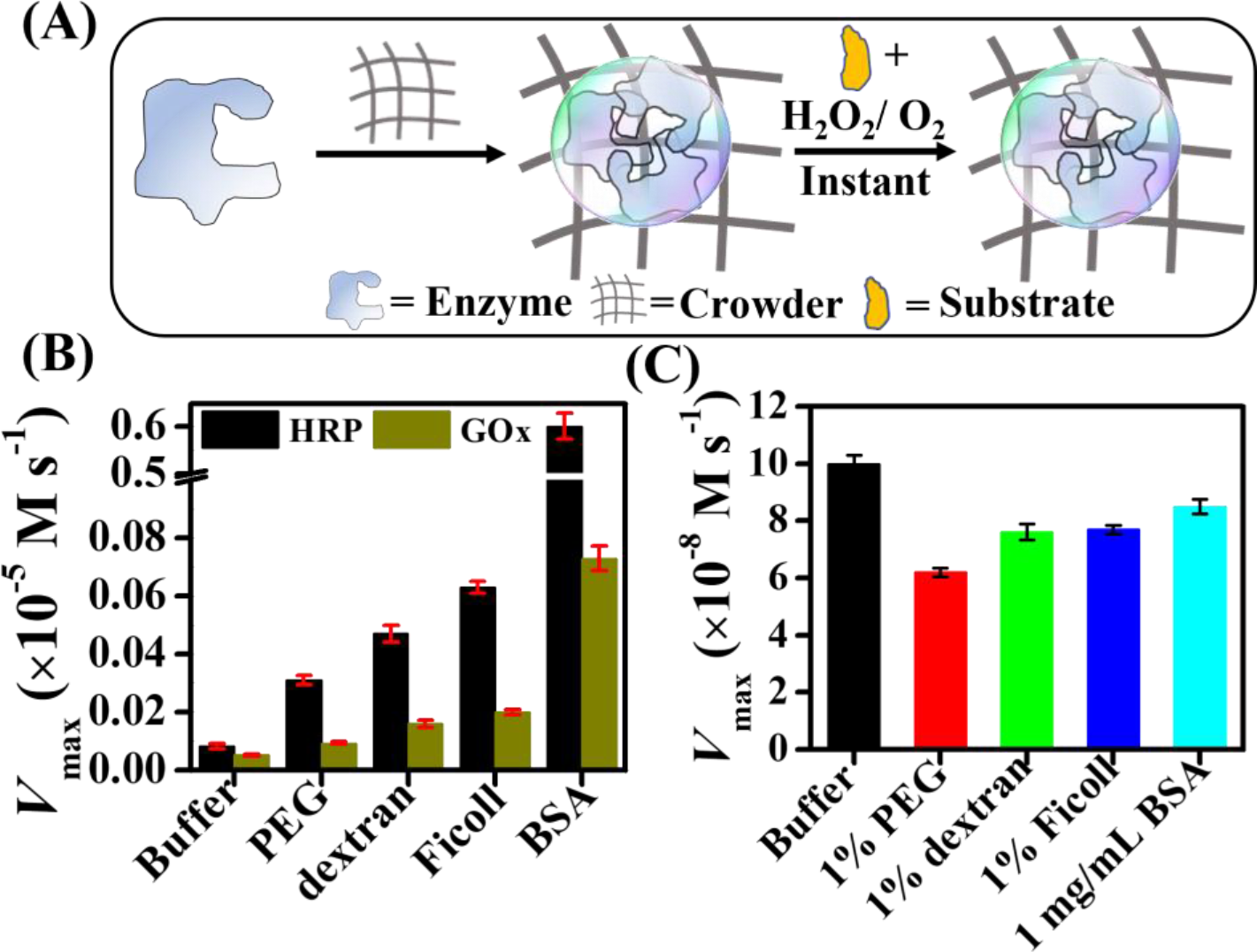
(A) Schematic illustration showing the enzymatic reaction after the LLPS in the presence of crowder. (B) The estimated *V*_max_ values of HRP and GOx in the absence and presence of 10% PEG, 10% dextran, 12.5% Ficoll, and 20 mg/mL BSA. (C) The estimated *V*_max_ values of HRP in the presence of 1% PEG, 1% dextran, 1% Ficoll, and 1 mg/mL BSA. The data represent the mean ± s.e.m. for three (n = 3) independent experiments.

The absorbance at 650 nm was monitored as a function of reaction time in the absence and presence of crowders (Figure S18). Reactions with phase separated droplets of HRP exhibit crowder-dependent variation in the rate. Interestingly, the enzymatic rate increases appreciably compared to that in the aqueous buffer for all the crowders (Figure S19). The observed enhancement is crowder dependent and maximum enhancement has been observed in the presence of 20 mg/mL BSA, whereas polymeric crowders show comparatively lower extent of enhancement. Similar enhancement in the enzymatic rates of phase-separated GOx has been observed in the presence of different crowders (Figures S20 and S21). The estimated *V*_max_ values are compared in Figure 6B. It is evident that for both HRP and GOx, the *V*_max_ increases in the presence of crowders relative to that in the aqueous buffer and more so for protein crowder. The *V*_max_ value of HRP and GOx increases by a factor of 62- and 14.3-times, respectively in the presence of BSA relative to that in the aqueous buffer. However, much lower extent (2.0–6.4-fold for HRP and 1.8–4.0-fold for GOx) of enhancement has been observed in the presence of polymeric crowders. The observed enhancement in the enzymatic rates could be due to the formation of phase separated droplet in the presence of crowders. In order to establish this argument, we performed enzymatic assays under the same experimental conditions with lower concentrations of crowders where no droplet formation was observed earlier. Control experiments reveal decrease in the *V*_max_ values compared to that in the aqueous buffer in the presence of 1% PEG, 1% dextran, 1% Ficoll, and 1mg/mL BSA (Figure 6C). Therefore, the unusual enhancement in the enzymatic rates of HRP and GOx arises primarily due to the formation of phase-separated droplets in the presence of crowders. On the other hand, the lower rates observed in the presence of polymeric crowder for both enzymes compared to that in protein crowder could be due to slow diffusion and lower extent of substrate partitioning in the viscous polymeric solutions. To establish this possibility, we performed time-dependent kinetic assays with phase-separated HRP in the presence of different crowders.

It should be noted that the reaction catalyzed by HRP provides a unique opportunity to test the effect of diffusion and substrate partitioning into the phase-separated droplets in a time-dependent manner as the reaction requires additional oxidant, H_2_O_2_, which is essential to initiate the catalytic transformation (Figure 7A).

**Figure 7.**
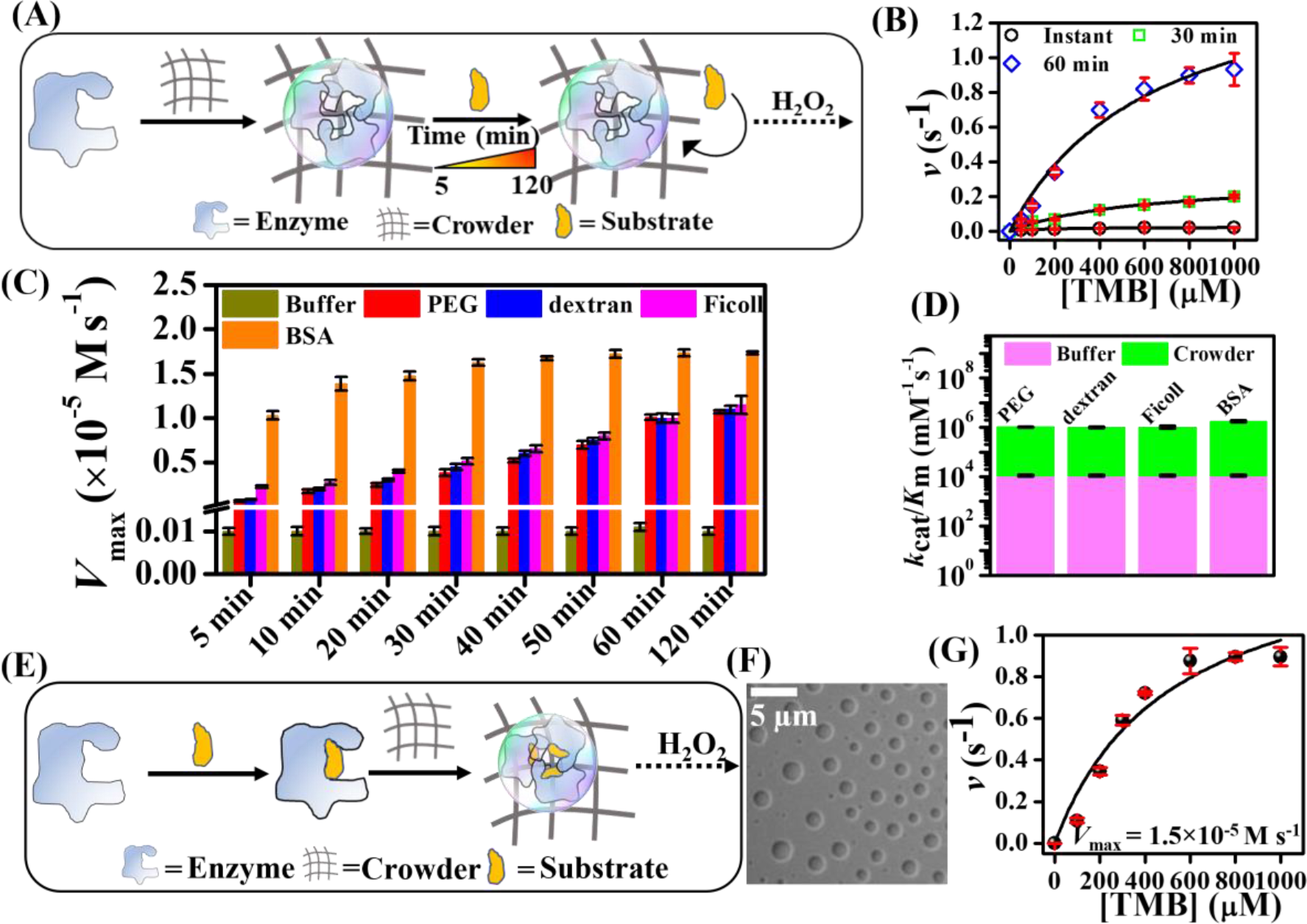
(A) Illustration of enzymatic reaction after the LLPS of enzyme in a time-dependent manner. (B) Michaelis-Menten plots of HRP as a function of TMB concentrations at different time intervals after the LLPS in 10% PEG. (C) The estimated *V*_max_ values of HRP in the absence and presence of various crowders at different time-intervals after the LLPS. (D) Plot of *k*_cat_/*K*_m_ values of HRP in buffer and different crowders. (E) Schematic illustration of enzymatic reaction after the LLPS of substrate-bound enzyme in the presence of 10% PEG. (F) Confocal DIC image of TMB-bound HRP in the presence of 10% PEG. (G) Michaelis-Menten plot after the LLPS of TMB-bound HRP. The data represent the mean ± s.e.m. for three (n = 3) independent experiments.

HRP catalyzed reactions were followed after the addition of TMB and initiated just after the addition of 8.8 mM H_2_O_2_ at a desire time interval (5–120 min). Surprisingly, the enzymatic rate of HRP in the presence of 10% PEG increases remarkably in a time-dependent manner as revealed from the Michaelis-Menten plots (Figure 7B). Similar time-dependent enhancement in the enzymatic rate has also been observed in the presence of dextran, Ficoll, and BSA as crowders (Figures S22–S24). Reaction monitored after 5 min of incubation of TMB with phase-separated droplets shows 7-, 8-, 22-, and 100-fold enhancement in the *V*_max_ value relative to that in the aqueous buffer in the presence of PEG, dextran, Ficoll, and BSA, respectively (Figure 7C). While a gradual increase in the *V*_max_ value has been observed in the presence of different crowders in a time-dependent manner, the *V*_max_ of HRP in the aqueous buffer under the same experimental conditions remains constant at 1 × 10^−7^ Ms^−1^. Notably, the kinetics of HRP in 20 mg/mL BSA differs from those in polymeric crowders in two aspects. First, a sharp increase in the *V*_max_ value has been observed for reaction monitored instantly after the addition of TMB and H_2_O_2_ in the presence of 20 mg/mL BSA. Second, the *V*_max_ of HRP in 20 mg/mL BSA saturates faster (within 30 min) than that in other polymeric crowders (within 60 min). Similar enhancement in the enzymatic rate of HRP has also been observed with OPD and ABTS as substrates in the presence of 10% PEG upon 60 min of incubation (Figures S25 and S26). The saturated values of *k*_cat_ and *K*_m_ for TMB are summarized in Table 2. It is evident that the *K*_m_ of TMB in the presence of different crowders remains almost unchanged relative to that estimated in the aqueous buffer under similar experimental conditions. Remarkably, the *k*_cat_ for TMB increases by 164-fold in the presence of 20 mg/mL BSA. The saturated value of catalytic efficiency (*k*_cat_/*K*_m_) of HRP increases by 114-, 91-, 114-, and 205-fold relative to the bulk aqueous phase in the presence of 10% PEG, 10% dextran, 12.5% Ficoll, and 20 mg/mL BSA, respectively (Figure 7D).

**Table 2.**
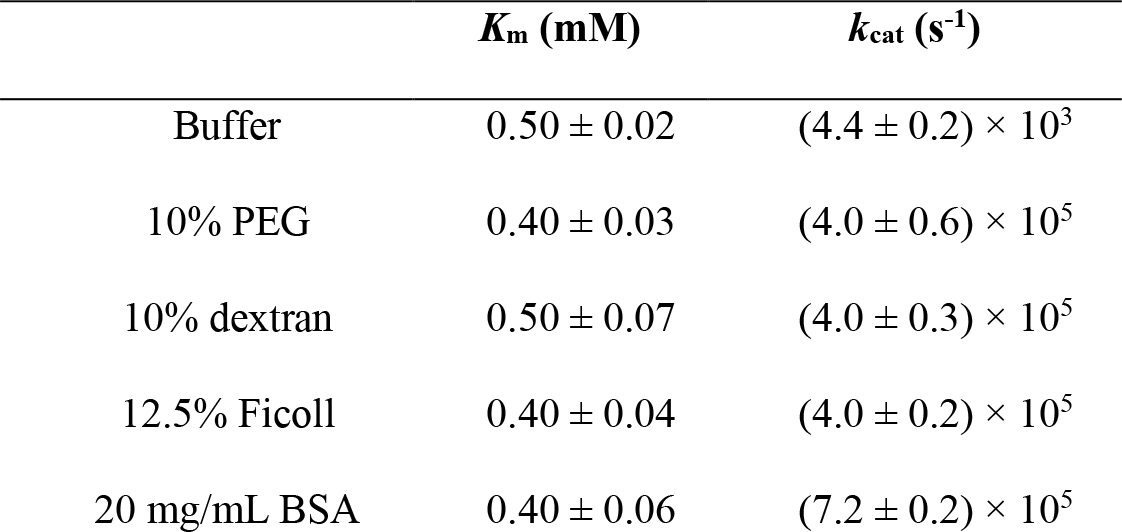
The Saturated Michaelis-Menten Parameters for HRP After the LLPS.

These findings suggest that LLPS has a tremendous influence on the enzymatic activity of HRP. In particular, the enhanced activity of phase-separated enzymes could be due to a combination of several factors. Among these, conformational change of the phase-separated enzyme is an important factor as revealed from our CD measurements. Second, the high local concentrations (mass action) of enzymes and substrates inside the phase-separated droplets can also accelerate the enzymatic kinetics. A control experiment with a 100-fold molar excess of HRP and substrates in the aqueous buffer reveals that the observed enhancement is not solely due to a simple mass action mechanism (Figure S27). Third, the confinement induced enhancement in the activities of bound water and substrates may also contribute positively to the overall enzymatic kinetics. Previously, it has been shown that confinement can significantly influence the feasibility, kinetics, and efficacy of a wide range of chemical transformations.^56,57^ Therefore, the remarkable enhancement in the catalytic efficiency of phase-separated enzymes originates due to a combined effect of altered conformation of the active enzymes, mass action, and confinement induced enhanced activities of water and substrates inside the phase separated droplets.

The observed differences in the kinetic parameters for polymeric and protein crowders clearly indicate the critical role of diffusion and partitioning of TMB inside the phase-separated droplets in crowded polymeric solutions. The aqueous polymeric solutions being more viscous compared to the aqueous BSA solution provide additional diffusion barriers.^24^ Moreover, it has been reported previously that TMB can specifically bind with polymeric crowders via the hydrophobic interactions.^21^ Therefore, the lower enzymatic rates in polymeric crowders could also be due to specific hydrophobic interactions of TMB with polymers. To further substantiate these arguments, we performed a control experiment with TMB-bound HRP after its LLPS in the presence of crowders (Figure 7E). Importantly, TMB-bound HRP exhibits a similar kind of LLPS in the presence of 10% PEG as revealed from confocal DIC image (Figure 7F). A Kinetic experiment with TMB-bound HRP after the LLPS in 10% PEG reveals a saturated *V*_max_ value of 1.5 × 10^−5^ M s^−1^ (Figure 7G). This high value of *V*_max_ obtained instantly after the phase separation of TMB-bound HRP suggests that the time-dependent changes observed earlier in the HRP kinetics originate exclusively from the slow diffusion and substrate partitioning inside the droplets in a crowded environment. Taken together, our findings clearly indicate the critical role of diffusion and substrate partitioning on the enzymatic kinetics in a crowded milieu with phase-separated droplets.

Finally, we seek to address the feasibility of GOx/HRP cascade reaction after their respective phase separation in the presence of 10% PEG and 20 mg/mL BSA (Figure 8A). CLSM measurements reveal spontaneous coalescence between HRP and GOx droplets upon mixing their phase-separated solution in the presence of 10% PEG (Figure 8B).

**Figure 8.**
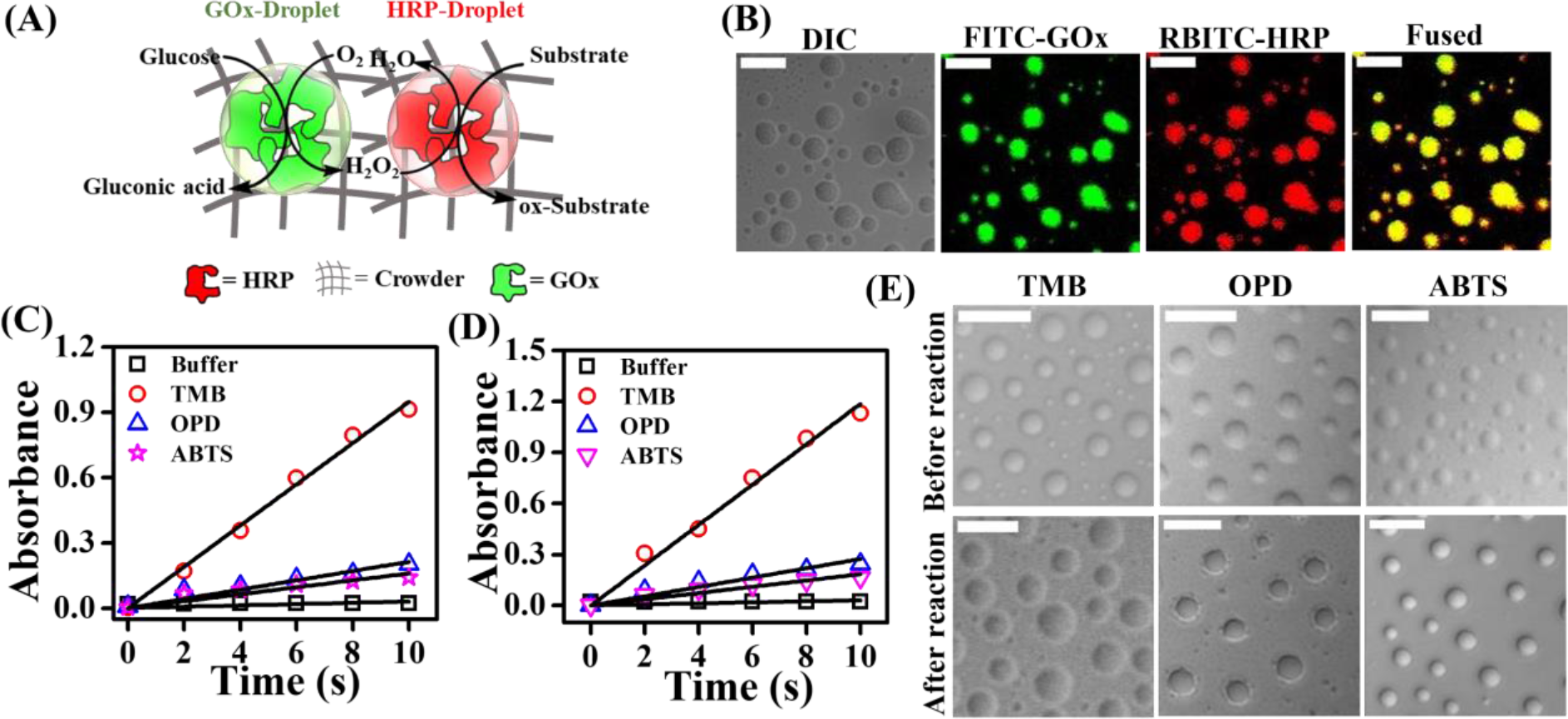
(A) Illustration of GOx/HRP cascade reaction in a crowded environment. (B) Confocal images showing the spontaneous coalescence between FITC-labeled GOx droplets and RBITC-labeled HRP droplets. Linearized plots of absorbance of oxidized substrates (TMB, OPD, and ABTS) for GOx/HRP cascade reactions in the absence and presence of (C) 10% PEG, and (D) 20 mg/mL BSA. (E) Confocal DIC images of droplets before and after the cascade reactions with different substrates. The scale bars correspond to 5 μm.

The uniform yellow signal appears exclusively from the interior of these droplets as a consequence of spontaneous fusion of red emitting RBITC-labeled HRP droplets and green emitting FITC-labeled GOx droplets. This observation can be explained by considering the isoelectric point (pI) of HRP (8.8) and GOx (4.2).^52^ It is expected that the droplet formed by HRP at pH 4.0 will have a net positive charge, whereas droplet formed by GOx at pH 7 will bear a net negative charge. Upon mixing, these oppositely charged droplets of GOx and HRP fuse spontaneously to yield droplets having both GOx and HRP. The cascade reaction was initiated after the addition of glucose into the reaction mixture. The reactions were monitored by observing the time-dependent formation of oxidized products of TMB, OPD, and ABTS. In buffer, the apparent rate constant (*k*_app_) of the cascade reaction with TMB as substrate is estimated to be 3.0 × 10^−3^ s^−1^, which increases to 9.5 × 10^−2^ and 1.2 × 10^−1^ s^−1^ in the presence of 10% PEG and 20 mg/mL BSA, respectively (Figure 8C, D). This 32-, and 40-fold enhancement in the *k*_app_ value of the GOx/HRP cascade reaction in the presence of crowders is unprecedented in the literature and indicates the active role of phase-separated droplets on the cascade activity in a crowded milieu. Notably, much lower extent of enhancement has been observed for OPD (7-, and 9-fold) and ABTS (5.3-, and 6-fold) as substrates in the presence of both PEG and BSA (Figure 8C, D). This contrasting behavior of substrate specificity could be either due to substrate-dependent alteration of the stability of fused droplets or due to the intrinsic physicochemical properties of substrates. To know the stability of GOx/HRP fused droplets in the presence of different substrates, we recorded the DIC images fused droplets before and after the cascade reactions (Figure 8E). It is evident that the morphologies of fused droplets remain unaltered after the cascade reactions, indicating that the droplets are stable in the presence of different substrates during the reactions. In contrast, the physicochemical properties of these substrates differ significantly from each other. The substrate TMB, being more hydrophobic than OPD and ABTS,^21^ partitioned preferentially inside the droplets containing both HRP and GOx. In addition, the hydrophobicity of fused droplets containing both GOx and HRP is comparatively higher than the individual droplets of GOx and HRP. On the other hand, hydrophilic substrates OPD and ABTS prefer to remain in the bulk aqueous phase, which lowers their partition coefficient. Therefore, our findings reveal that the physicochemical properties of phase-separated droplets and substrates dictate the specificity and selectivity of the enzymatic cascade reactions under heterogeneous and crowded environments. Finally, control experiments with two other enzymes, namely trypsin, and alcohol dehydrogenase reveal a similar crowding induced spontaneous LLPS phenomenon (Figure S28), suggesting the generality of this physiologically relevant unique event in regulating the efficacy of complex biocatalytic reactions under heterogeneous and crowded environments.

Based on our findings, we propose a mechanistic model where enzymatic transformations can take place via three main competing pathways in a crowded environment (Scheme 2). We believe that spatio-temporal regulation of these pathways determines the fate of enzymatic transformations in a crowded environment. In pathway 1, enzymes bind with substrates in a diffusion-controlled manner under the crowded environment and subsequently transform into products. Most of the earlier reported enzymatic kinetics follow this pathway, where reactions were monitored just after the addition of enzymes into the reaction mixture in the presence of crowders. The enzymatic activity in pathway 1 decreases significantly compared to that in the aqueous buffer due to the diffusion barriers in a crowded environment. The present study is first of its kind to discover the presence of two other competing pathways in the presence of macromolecular crowders. In pathway 2, enzymes first undergo spontaneous LLPS in the presence of crowders via the formation of liquid-like droplets (Scheme 2). Subsequently, substrate binding via time-dependent partitioning into these phase-separated droplets leads to enzyme-substrate adducts. These substrate-bound phase-separated enzymes ultimately undergo catalytic transformations in the presence of oxidants.

**Scheme 2.**
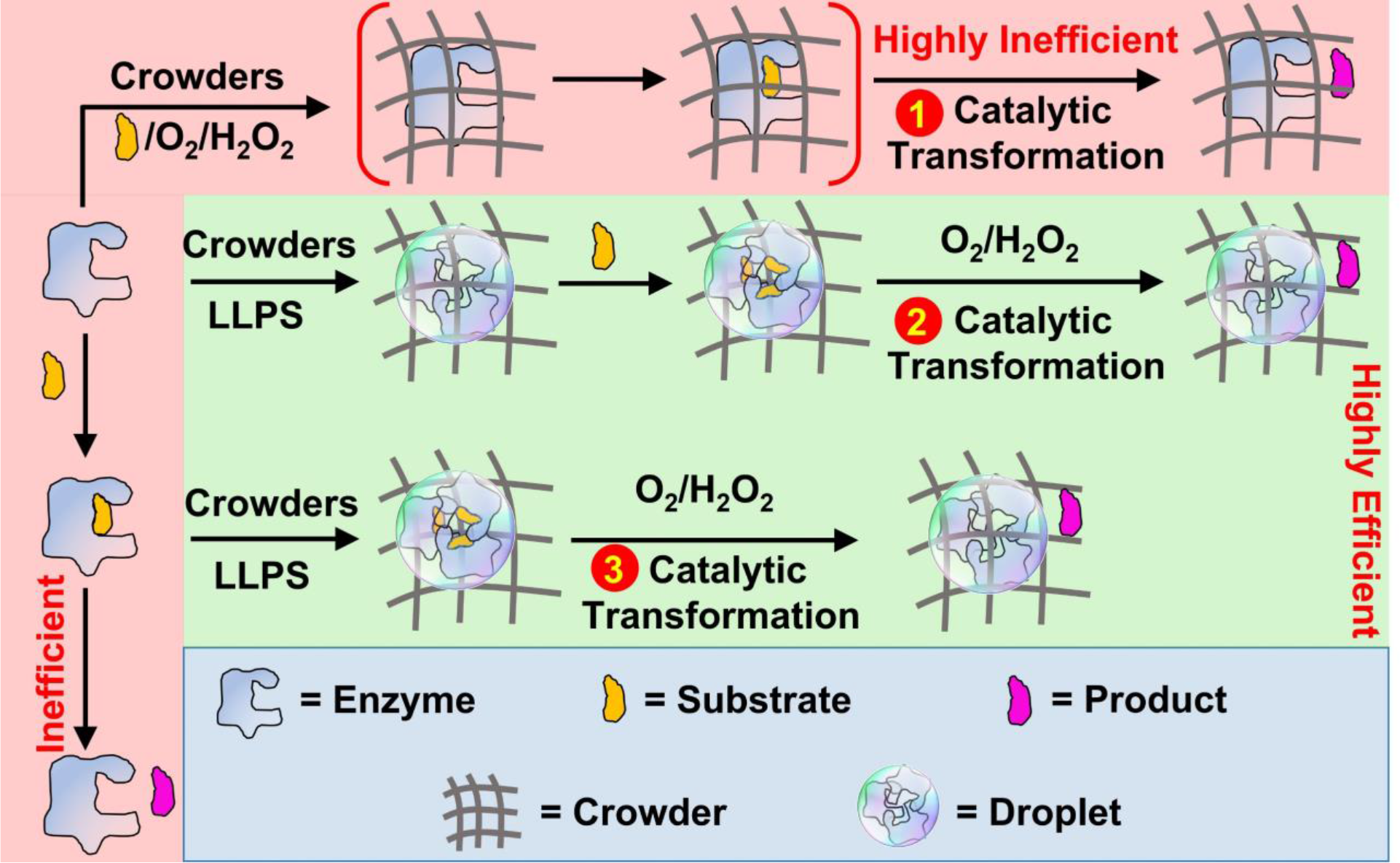
Illustration of Competing Pathways for Enzymatic Reactions under the Crowded Environment.

The enzymatic activity enhances appreciably in pathway 2 in a time-dependent manner due to the slow diffusion and partitioning of substrates into the phase-separated droplets in the presence of crowders. Biomolecules can interact with neighboring biomolecules via various noncovalent interactions including electrostatic, hydrophobic, hydrogen bonding, aromatic, and cation-π interactions. In particular, we have shown that both HRP and GOx undergo homotypic LLPS in the presence of inert crowders via combination of multivalent electrostatic and hydrophobic interactions. These phase-separated droplets contain conformationally altered active enzymes. In pathway 3, substrate-bound enzymes first undergo LLPS in the presence of crowders to yield liquid-like droplets. These phase-separated substrate-bound enzymes take part in the catalytic transformations in the presence of oxidants without any diffusion and partitioning barriers for substrates. Pathway 3 shows maximum enhancement in the enzymatic activity in the presence of crowders relative to that in the aqueous buffer and other pathways (Scheme 2). Therefore, our present study is one of its kind to illustrate the complex interplay of LLPS and enzymatic transformations under the crowded environment. While significant hindrance in the enzymatic activity has been observed before the LLPS of enzymes, remarkable enhancement has been observed in the catalytic activity after the LLPS of enzymes. The enhanced activity is due to a combined effect of conformational alteration of active enzymes, mass action, and confinement-induced alteration in the activities of water and substrates. In the physiological context, we believe that Nature must utilize this spatio-temporal synchronization of LLPS, substrate binding, and enzymatic transformation in a highly regulated manner to tune the efficiency of various parallel and tandem complex metabolic reactions under the highly crowded environments.

## 3. Conclusions

In summary, we have discovered a unique and unprecedented phenomenon of LLPS of biologically active enzymes under the crowded environments. The presence of macromolecular crowders induces homotypic LLPS of HRP and GOx via the formation of liquid-like membraneless droplets. Droplet formation has been shown to be inhibited completely at high temperature of 90 °C for both enzymes, signifying enthalpically controlled UCST profiles. We have shown that the observed LLPS events of both these enzymes are mainly driven by electrostatic as well as hydrophobic intermolecular interactions. Using a range of kinetic experiments, we have illustrated the active role of phase-separated droplets behind the crowding induced enhancement of enzymatic activity under physiological conditions. Our mechanistic model represents the first proof of concept example of enhanced enzymatic activity and selectivity inside the phase-separated liquid-like droplets under a crowded milieu. The enhanced activities inside the phase-separated droplets have been explained by considering conformational change of the active enzymes, mass action, and confinement induced alteration of the activity of surrounding water and substrates. Our present study illustrates that precise synchronization of LLPS and catalytic transformation is crucial to realize the enhanced activity and selectivity inside the phase-separated droplets. The present discovery of phase separation-induced active regulation of enzymatic activity and selectivity opens the door of a new avenue in biocatalysis with a vast scope in industrial applications.

## ASSOCIATED CONTENT

### Supporting Information

Experimental details, characterization techniques, and Figures S1–S28.

## AUTHOR INFORMATION

### Notes

The authors declare no competing financial interests.

## Acknowledgements

The authors acknowledge Indian Institute of Technology Indore for providing financial support, infrastructure, and instrumentation facilities; SIC, IIT Indore, for instrumental facilities. B.S. acknowledges the Council of Scientific & Industrial Research (CSIR), India, for research fellowship.

