## Supplemental file for "Biomolecular Condensate Regulates Enzymatic Activity under Crowded Milieu: Synchronization of Liquid-Liquid Phase separation and Enzymatic Transformation"

AUTHOR INFORMATION

**Corresponding Author**

[*]

### Table of Contents


| S.No. | | Page No. |
|---|---|---|
| 1 | Materials | S3 |
| 2 | Supporting Methods | S3-S7 |
| 3 | Characterization Techniques | S7-S8 |
| 4 | Supporting Figures | S9-S36 |
| 5 | References | S37 |

#### 1. Materials

Horseradish peroxidase (HRP), glucose oxidase (GOx), trypsin, yeast alcohol dehydrogenase, polyethylene glycol (PEG 8000), Ficoll 400, dextran 70, methoxy polyethylene glycol amine (mPEG-$NH_2$ 5000), glucose, sodium thiocyanate (NaSCN), 1,6-hexanediol, rhodamine B isothiocyanate (RBITC), fluorescein-5-isothiocyanate (FITC), bovine serum albumin (BSA, ≥ 99%, essentially fatty acid-free), sodium citrate tribasic dihydrate, potassium chloride, hydrochloride acid, Tris buffer, Hellmanex III, and the Pur-A-Lyzert dialysis kit (molecular weight cutoff 3.5 kDa) were purchased from Sigma-Aldrich. 3,3',5,5'-tetramethylbenidine (TMB), o-phenylenediamine (OPD), and 30% hydrogen peroxide ($H_2O_2$) were purchased from Loba Chemie. 2,2'-Azinobis [3-ethylbenothiazoline-6-sulphonic acid]-diammonium salt (ABTS) was purchased from TCI. Sodium dihydrogen phosphate monohydrate ($NaH_2PO_4.H_2O$), di-sodium hydrogen phosphate heptahydrate ($Na_2HPO_4.7H_2O$), sodium acetate trihydrate ($CH_3COONa.3H_2O$), acetic acid ($CH_3COOH$), sodium bicarbonate ($NaHCO_3$), sodium carbonate anhydrous ($Na_2CO_3$), sodium chloride (NaCl), and dimethyl sulfoxide (DMSO) were purchased from Merck. Milli-Q water was obtained from a Millipore water purifier system (Milli-Q integral).

#### 2. Methods

##### Predicting LCDs and IDRs of HRP and GOx

To predict the low complexity domains (LCDs) and disorder regions (IDRs) in HRP and GOx, we used Simple Molecular Architecture Research Tool (SMART) (http://smart.embl-heidelberg.de/) and IUPred2, (https://iupred2a.elte.hu/) respectively. IUPred2 data were then plotted using the OriginPro 8.1 software.

**Preparation of Buffer Solutions**

Different buffer solutions with pH values 2.0, 4.0, 7.4, 9.0, and 10.0 were prepared by using Milli-Q water. The strength of each buffer solution was kept fixed at 10 mM. Hydrochloride acid-potassium chloride buffer (pH 2.0), sodium acetate buffer (pH 4.0), phosphate buffer (pH 7.4), tris buffer (pH 9.0), and carbonate-bicarbonate buffer (pH 10.0) were used individually in the presence of 50 mM NaCl.

**Preparation of Crowder and Substrate Solutions**

10% (w/v) PEG, 10% (w/v) dextran, 12.5% (w/v) Ficoll were prepared from a stock solution of 40% (w/v) PEG 8000, 40% (w/v) dextran 70, and 40% (w/v) Ficoll 400, respectively. 20 mg/mL BSA was prepared from the stock solution of 332 mg/mL BSA. Stock solution of 20 mM TMB was prepared in DMSO, while the stock solutions of 20 mM OPD and ABTS were prepared in Milli-Q water. The stock solutions of these substrates were used immediately after preparation.

**Labeling of Enzymes and Crowders with Fluorescent Dyes**

The concentration of enzymes was estimated spectrophotometrically using the reported extinction coefficient of $1.02 \times 10^5$ $M^{-1}$ $cm^{-1}$ ($\lambda = 403$ nm) for HRP, $4.41 \times 10^4$ $M^{-1}$ $cm^{-1}$ ($\lambda = 280$ nm) for GOx, $8.80 \times 10^2$ $M^{-1}$ $cm^{-1}$ ($\lambda = 253$ nm) for trypsin, and $1.89 \times 10^5$ $M^{-1}$ $cm^{-1}$ ($\lambda = 280$ nm) for alcohol dehydrogenase. HRP and alcohol dehydrogenase were labeled with RBITC, whereas GOx and trypsin were labeled with FITC according to an earlier reported method.[1] In brief, 25 pM or 0.5 μM of each HRP and GOx were mixed with RBITC and FITC, respectively in a molar ratio of 1:10 ([Enzyme]: [Dye]). Similarly, 60 nM alcohol dehydrogenase and 42 nM trypsin were mixed with RBITC and FITC, respectively in a molar ratio of 1:10 ([Enzyme]:[Dye]). These enzyme-dye mixtures were incubated for 4 h at room temperature followed by 6 h at 4 °C with constant stirring (500 rpm). After completion of the reaction, the excess dye was removed by dialysis (molecular weight cutoff 3.5 kDa) against 10 mM PBS at 4 °C for 12 h. Finally, the labeled enzymes were stored at 4 °C. The estimated labeling efficiency of HRP and GOx is found to be 77.3% and 84.1%, respectively. The same procedure was followed for the labeling of mPEG-$NH_2$ and BSA with RBITC.

**Phase Separation Assays in the Presence of Crowders**

0.5 μM HRP and 0.5 μM GOx were incubated individually with 10% PEG, 12.5% Ficoll 400, 10% dextran 70, and 20 mg/mL BSA at 37 °C for 1 h. The droplet formation was monitored using CLSM with 100%-labeled enzymes and field-emission scanning electron microscopy (FESEM). The effect of incubation time on droplet formation was monitored by varying the equilibration time (5 min,15 min, 30 min, and 60 min) after the mixing of 10% PEG with the fluorescently labeled HRP and GOx. The droplet formation in the presence of 10% PEG was monitored under different conditions by varying temperature (4, 37, 70, 80, and 90 °C), pH (4.0, 7.4, 9.0, and 10.0), NaCl concentration (50 mM, 500 mM, 1000 mM, 2000 mM, and 3000 mM), NaSCN concentration (0.5 M, 1 M, 2 M, and 3 M), and 1,6-hexanediol concentration (1%, 3%, 6%, and 10%) upon 1 h of incubation.

**Enzymatic Assays**

**pH-Dependent Enzymatic Activity in Aqueous Buffer**

The concentration of HRP and GOx was kept fixed at 25 pM for all the kinetic experiments. All the enzymatic assays were performed in triplicates (n=3) at 37 °C. The optimum activity of HRP and GOx was estimated in aqueous buffer as a function of pH in the range of 2–10 at 37 °C with TMB (0–1000 μM) and glucose (0–300 mM) as substrate, respectively. HRP catalyzed reactions were monitored using UV-vis spectrophotometer immediately after the addition of 8.8 mM $H_2O_2$ via recording the absorbance of oxidized TMB at 650 nm. GOx activity was measured according to the earlier report.[2] After purging $O_2$ gas for 15 min into the reaction mixtures, the kinetics were monitored immediately after the addition of different concentrations of glucose. The final product gluconic acid was assayed by reaction with hydroxylamine and subsequent complexation with $Fe^{3+}$, which led to a red color complex hydroxamate-$Fe^{3+}$ ($\lambda_{max}$ = 505 nm).

**Enzymatic Assays before Phase Separation in the presence of Crowders**

Enzymatic kinetics before the phase separation were followed instantly after the addition of enzymes into the aqueous solution of substrate in the presence of different crowders (10% PEG, 10% dextran, 12.5% Ficoll, 20 mg/mL BSA). The enzymatic kinetics of HRP and GOx were monitored at their optimum pH values of 4.0 and 7.4, respectively. All the data points were reported as mean ± s.e.m. Statistical analyses were performed via a two-tailed, unpaired Student's *t*-test with $^{***}P$ value <0.001, $^{**}P$ value <0.01 and not significant (NS; $P$ >0.05) using Excel software.

**Enzymatic Assays After Phase Separation**

Enzymatic assays after the phase separation were performed in three different sets. In the first set, enzymes were allowed to undergo phase separation in the presence of different crowders for 1 h. Subsequently, HRP and GOx kinetics were monitored instantly after the addition of respective substrates. In the second set, the kinetics of phase separated HRP were monitored after the addition of TMB (0–1000 μM) in a time-dependent manner. Reactions were initiated after the addition of 8.8 mM $H_2O_2$ at a definite time interval (5–120 min). The enzymatic activity of HRP in the presence of 10% PEG was also monitored with OPD (0-1000 μM) and ABTS (0-1000 μM) in pH 4.0 acetate buffer via recording the absorbance at 420 and 450 nm, respectively. In the third set, HRP was first allowed to bind with TMB (0–1000 μM) in pH 4.0 aqueous buffer at 37 °C for 30 min. Subsequently, substrate bound HRP was mixed with 10% PEG and incubated further for 15 min at 37 °C. Finally, reaction was initiated after the addition of 8.8 mM $H_2O_2$ into the reaction mixture.

**Estimation of Michaelis-Menten Parameters**

The absorbance values obtained from UV-vis spectrometer were plotted as a function of time using OriginPro 8.1 software and then the data were linearly fitted for each concentration. The slopes corresponding to these fitted graphs were plotted against substrate concentrations. These data were finally fitted with the Michaelis-Menten equation according to the following expression,

$$V_0 = \frac{V_{max}\,[S]}{K_m + [S]} \quad (1)$$

where, $V_0$ is the initial velocity, [S] is the molar concentration of substrate, $V_{\mathrm{max}}$ is the maximum velocity, and $K_{\mathrm{m}}$ is the Michaelis-Menten constant. We used non-linear curve fit analysis in OriginPro 8.1 software to estimate the fitted parameters $V_{\mathrm{max}}$, and $K_{\mathrm{m}}$. Finally, $k_{\mathrm{cat}}$ was calculated by dividing the $V_{\mathrm{max}}$ by the total enzyme concentration used.

**Enzymatic Cascade Reaction**

25 pM HRP and 25 pM GOx were incubated individually with either 10% PEG or 20 mg/mL BSA for 1 h at 37 °C in pH 4.0 acetate buffer and pH 7.4 phosphate buffer, respectively. Subsequently, both the solutions were mixed by taking equal volumes from each solution. Next, 1 mM of a substrate either TMB, OPD, or ABTS was added and further allowed to equilibrate for 1 h at 37 °C. Afterward, the reaction mixture was purged with $O_2$ gas for 15 min. Finally, the absorbance values of oxidized substrates were recorded immediately after the addition of 1 mM Glucose. Similarly, control experiments without crowders were performed under the same experimental conditions.

#### 3. Characterization Techniques

##### UV-vis Spectroscopy

Absorption spectra and reaction kinetics were monitored by using PerkinElmer UV/VIS/NIR spectrometer in a quartz cuvette (1 cm × 1 cm).

##### Confocal Laser Scanning Microscopy (CLSM)

The CLSM images were captured with an inverted confocal microscope (Olympus fluoView, model FV1200MPE, IX-83) using an oil immersion objective (100×1.4 NA). Diode laser sources at 488, and 559 nm were used to excite the samples by using appropriate dichroic and emission filters (green channel: 490-550 nm; and red channel: 560-650 nm) in the optical path. A 20 μL aliquot of the sample solution was drop-casted onto cleaned glass slides and sandwiched with Blue Star coverslip. Finally, the sides of the coverslips were sealed with commercially available nail paint, and then images were captured.

##### Field-Emission Scanning Electron Microscopy (FESEM)

The FESEM images were captured by using a Supra 55 Zeiss field-emission scanning electron microscope. For FESEM measurements, the dried samples were initially coated with gold.

##### Circular Dichroism (CD) Spectroscopy

CD spectra were recorded on a JASCO J-815 CD spectropolarimeter using a quartz cell of 1 mm path length and scans were recorded from 190-260 nm and 380-430 nm at 37°C with a slit width of 1 mm and a speed of 50 nm/min.

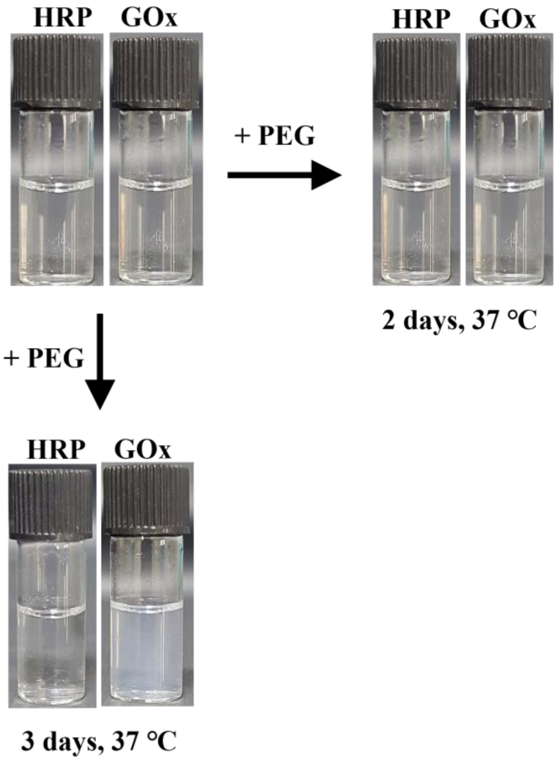


**Figure S1.** Daylight photographs of aqueous solutions of HRP and GOx in the absence and presence of 10% PEG in pH 4.0 acetate buffer.

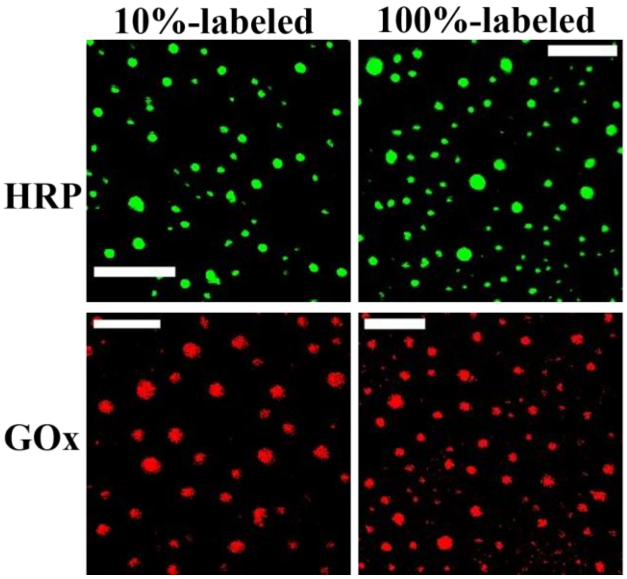


**Figure S2.** CLSM images of droplets in the presence of 10% PEG with 10%- and 100%-labeled enzymes. The scale bars correspond to 5 μm.

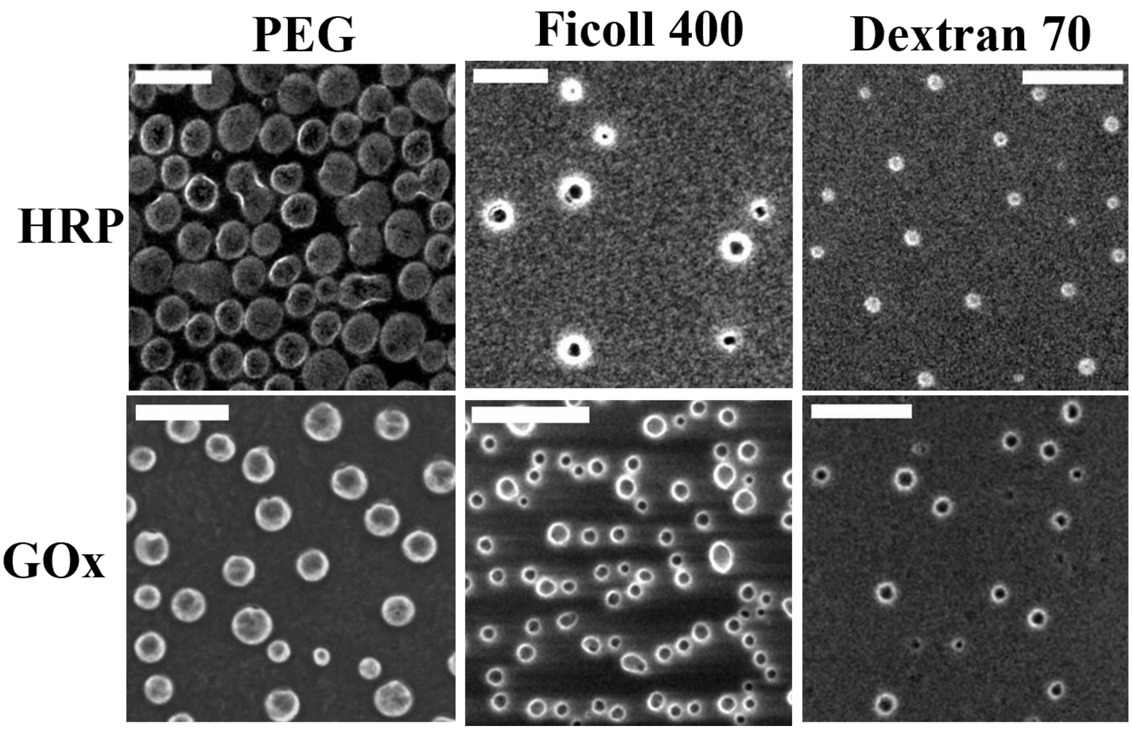


**Figure S3.** FESEM images of HRP and GOx droplets in the presence of polymeric crowders. The scale bars correspond to 5 μm.

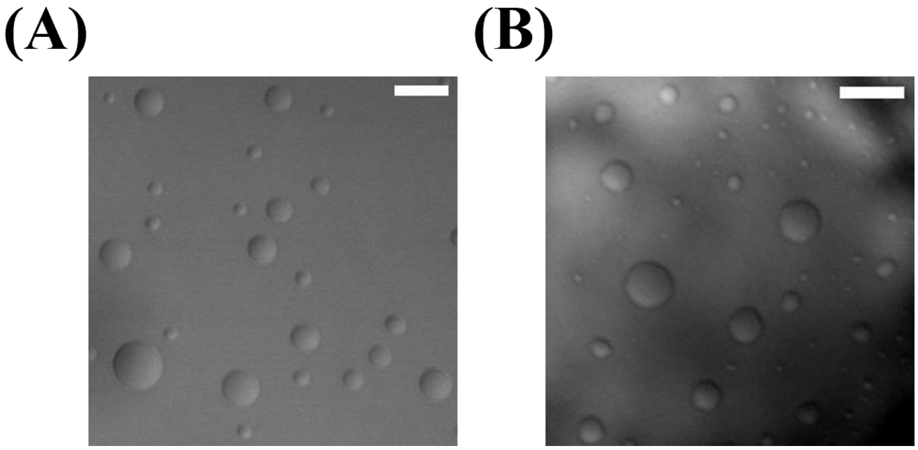


**Figure S4.** Confocal DIC images showing the stability of droplets (A) HRP, and (B) GOx over a period of 15 days. The scale bars correspond to 5 μm.

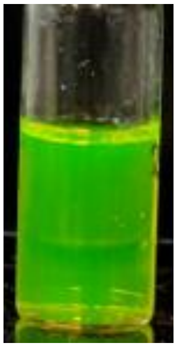

**Figure S5.** Daylight photograph of the isotropic mixture of PEG in aqueous buffer in the presence of FITC.

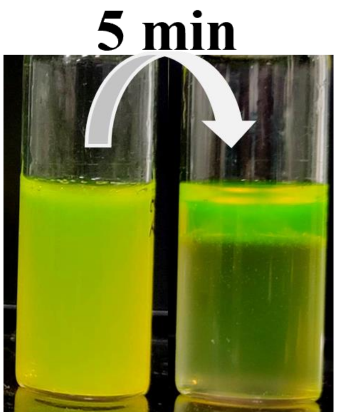


**Figure S6.** Daylight photographs of aqueous mixtures of PEG and citrate salt in the presence of FITC.

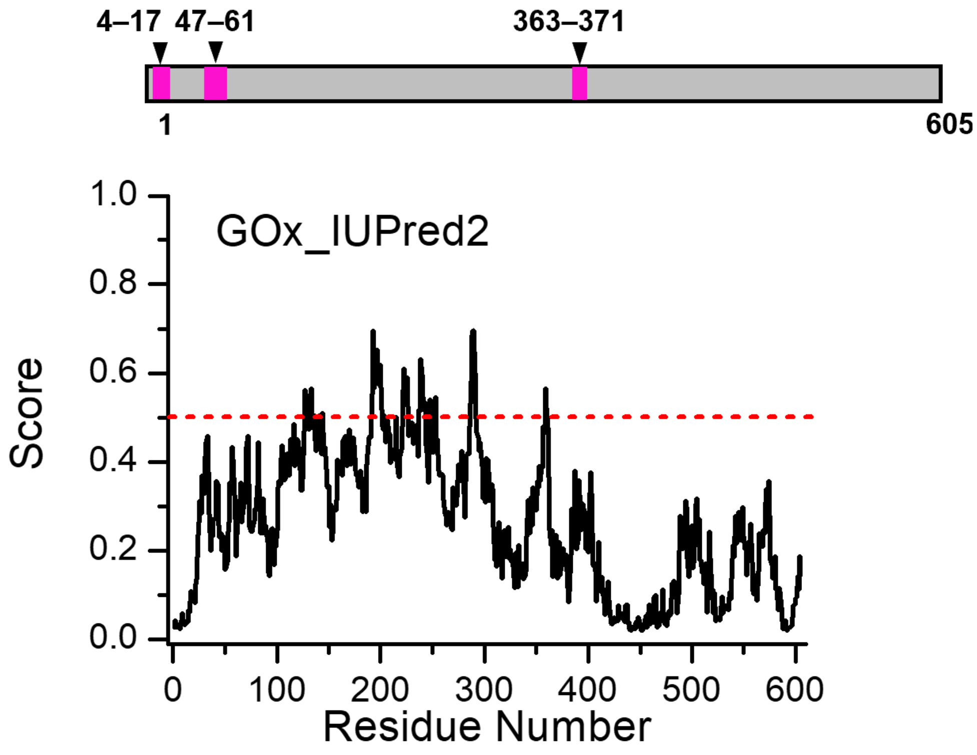


**Figure S7.** Predictive algorithm showing LCDs and IDRs of GOx.

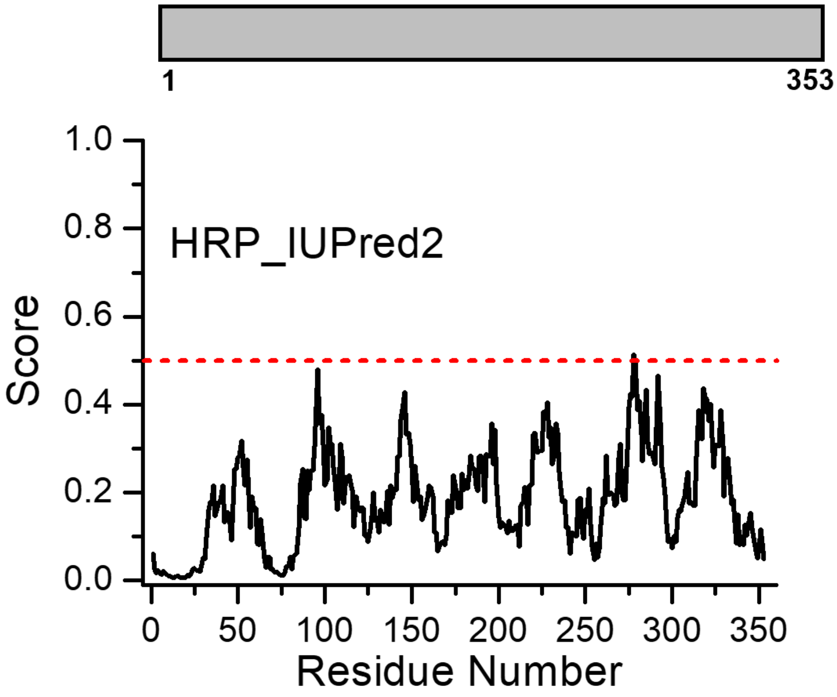


**Figure S8.** Predictive algorithm showing LCDs and IDRs of HRP.

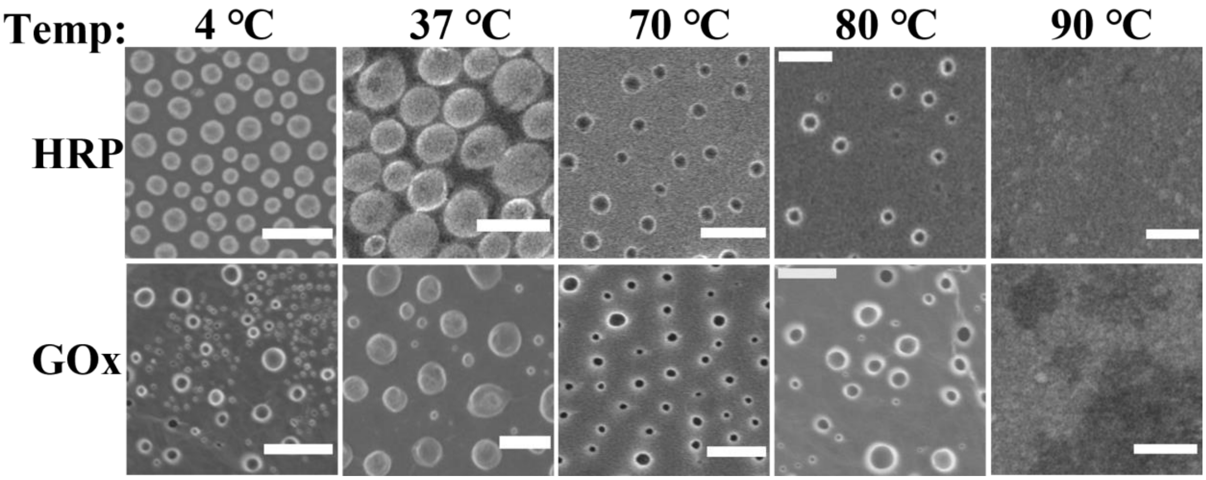


**Figure S9.** FESEM images showing the stability of HRP and GOx droplets as a function of temperature. The scale bars correspond to 5 μm.

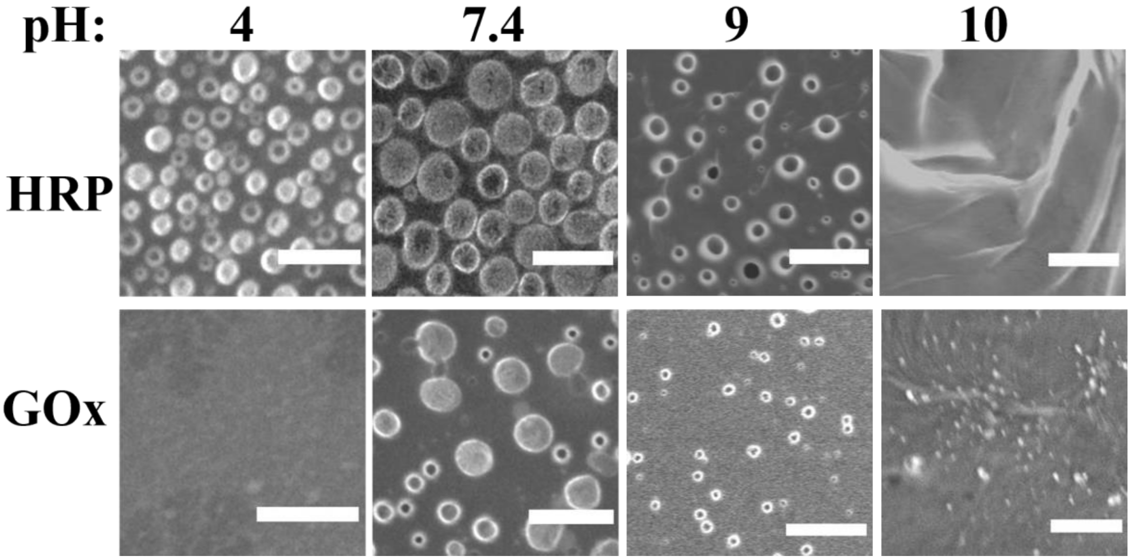


**Figure S10.** FESEM images showing the stability of HRP and GOx droplets as a function of pH. The scale bars correspond to 5 μm.

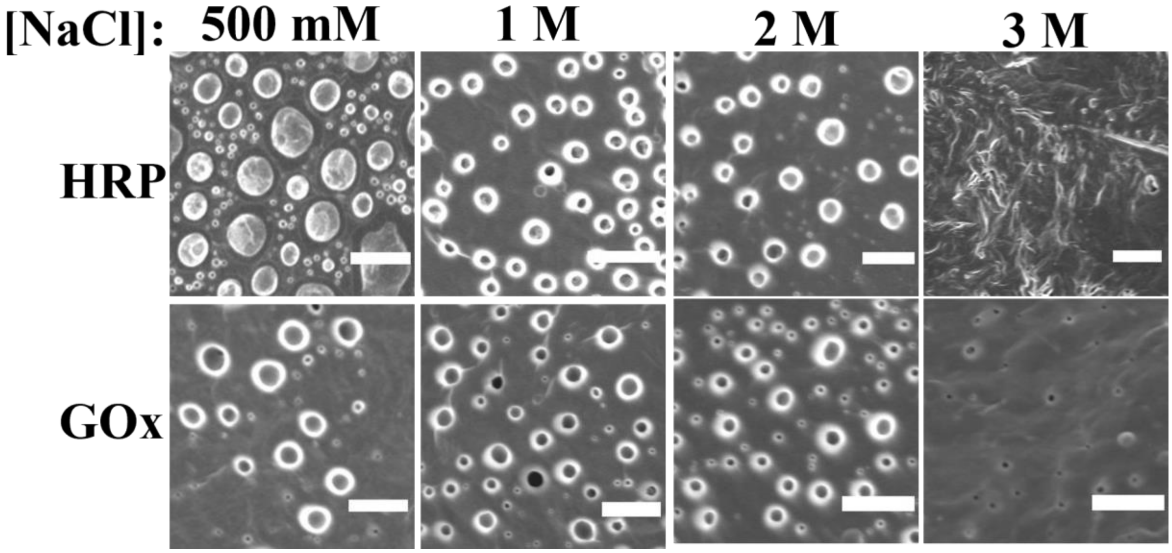


**Figure S11.** FESEM images showing the stability of HRP and GOx droplets as a function of NaCl concentrations. The scale bars correspond to 5 μm.

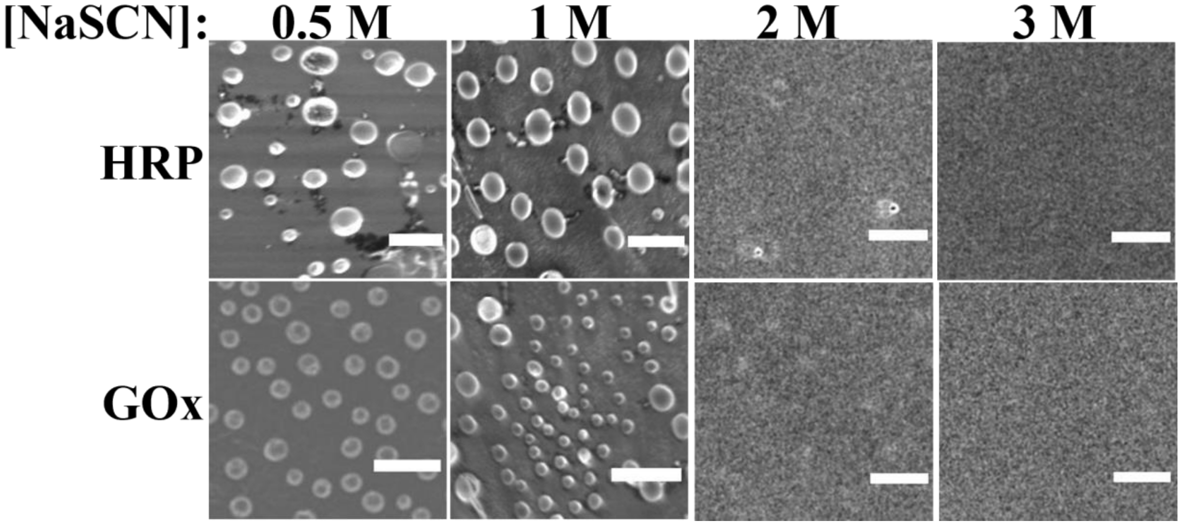


**Figure S12.** FESEM images showing the stability of HRP and GOx droplets as a function of NaSCN concentrations. The scale bars correspond to 5 μm.

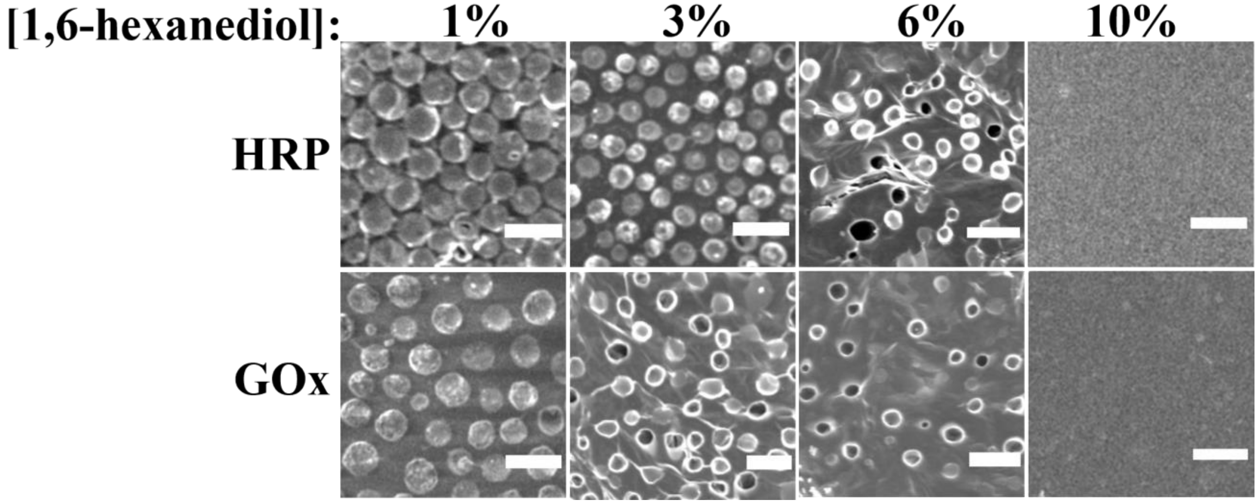


**Figure S13.** FESEM images showing the stability of HRP and GOx droplets as a function of 1,6-hexanediol concentrations. The scale bars correspond to 5 μm.

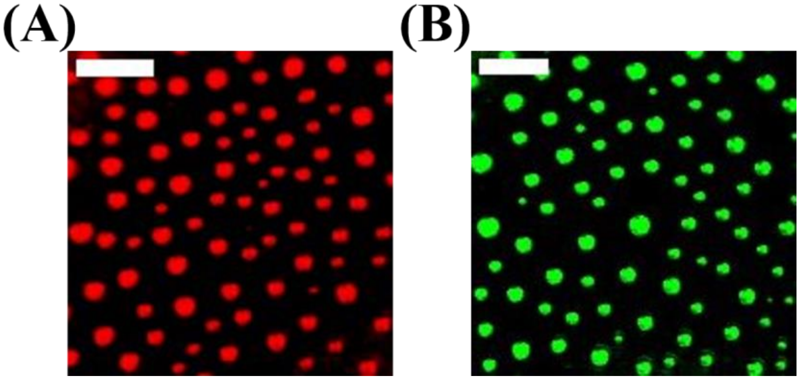

**Figure S14.** CLSM images of (A) RBITC-labeled HRP, and (B) FITC-labeled GOx in the presence of 10% PEG at 37 ℃ with an enzyme concentration of 25 pM. The scale bars correspond to 5 μm**.**

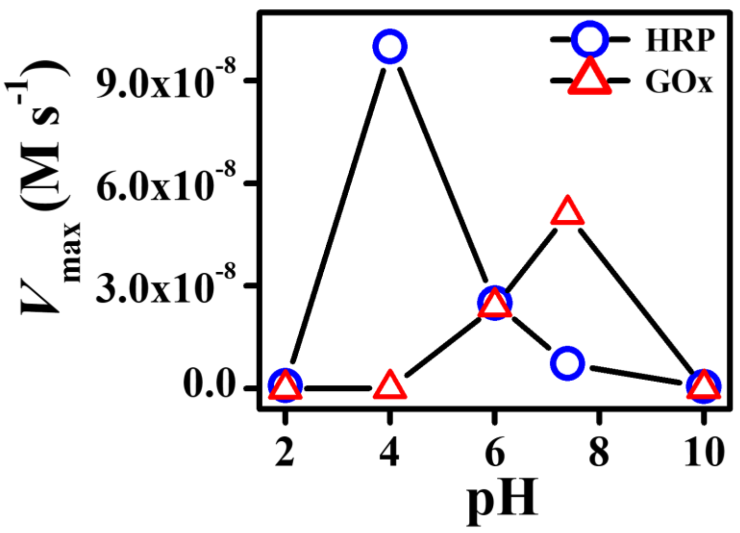


**Figure S15.** Plot of $V_{max}$ versus pH for HRP and GOx catalyzed reactions at 37 °C.

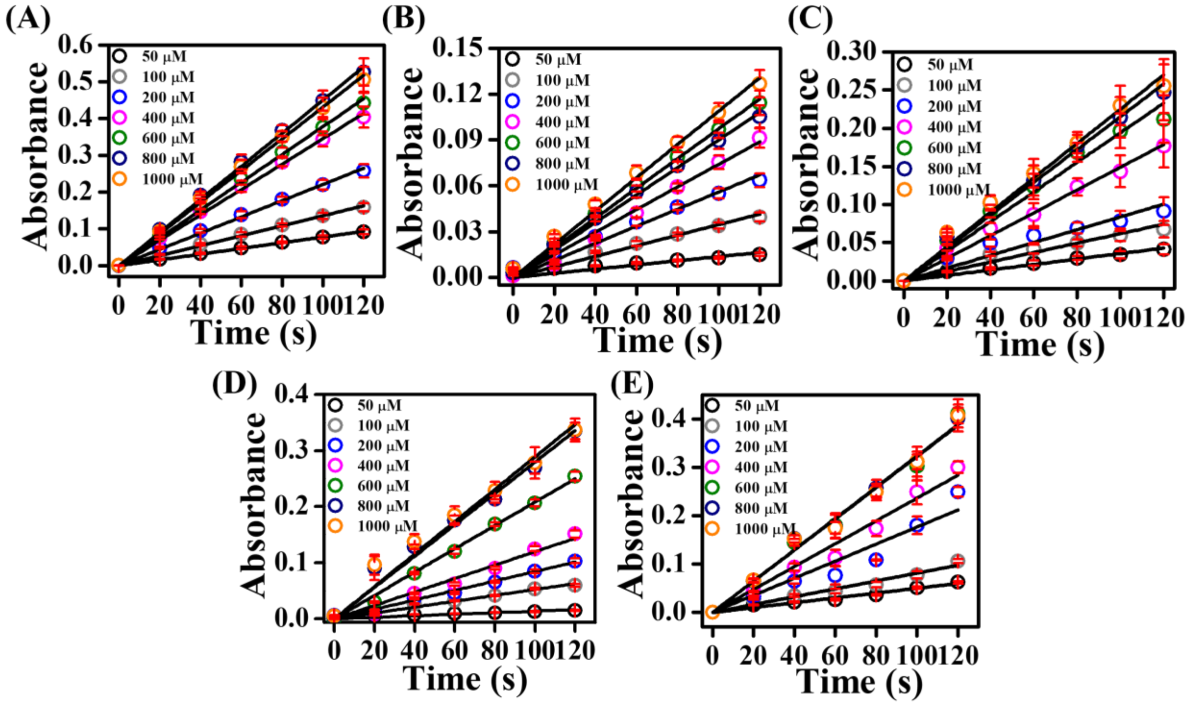


**Figure S16.** Plots of absorbance ($\lambda$ = 650 nm) versus time as a function of TMB concentrations in (A) Buffer, (B) 10% PEG, (C) 10% dextran, (D) 12.5% Ficoll, and (E) 20 mg/mL BSA in pH 4.0 acetate buffer at 37 °C.

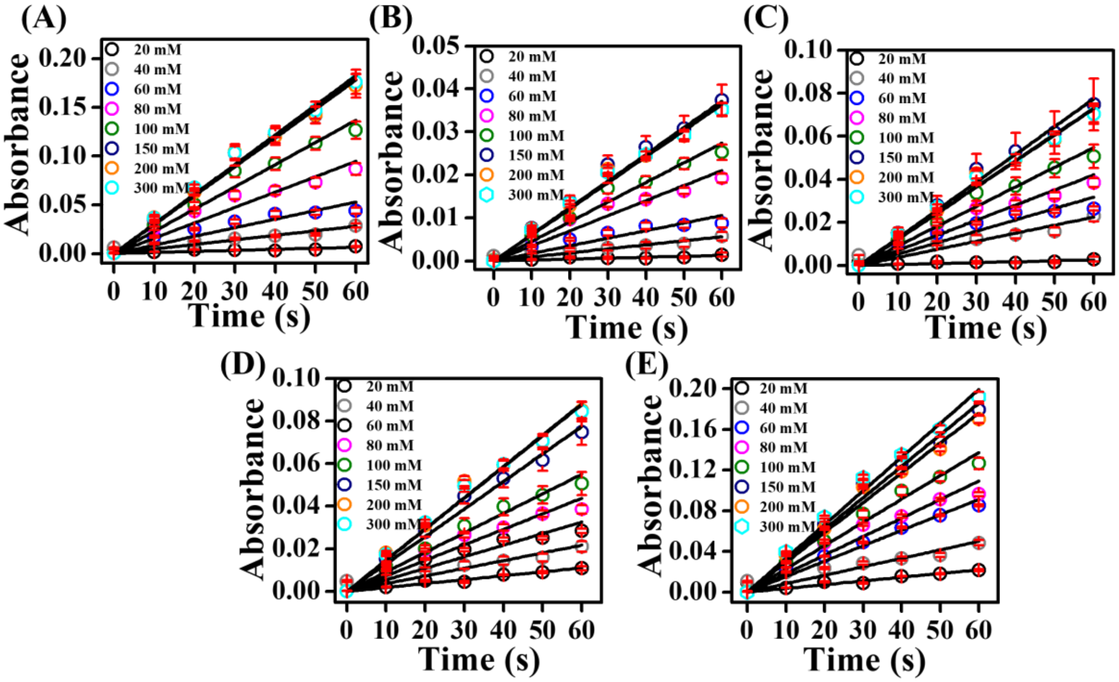


**Figure S17.** Plots of absorbance ($\lambda$ = 505 nm) versus time as a function of glucose concentrations in (A) Buffer, (B) 10% PEG, (C) 10% dextran, (D) 12.5% Ficoll, and (E) 20 mg/mL BSA in pH 7.4 PBS at 37 °C.

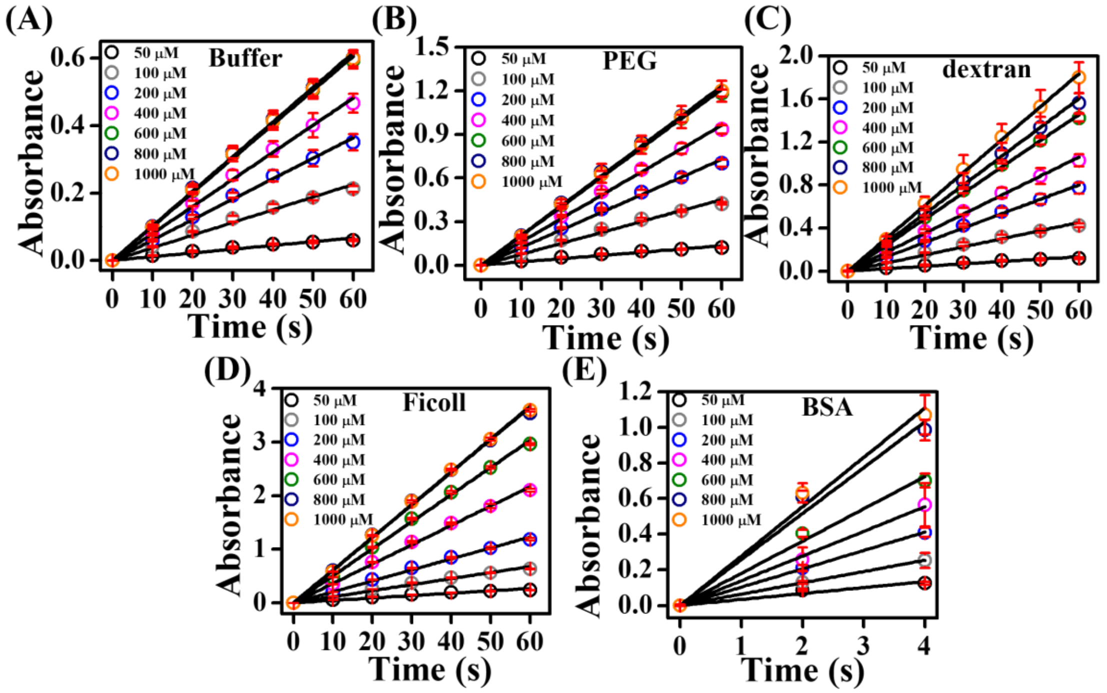


**Figure S18.** Plots of absorbance ($\lambda$ = 650 nm) versus time as a function of TMB concentrations in the absence and presence of different crowders in pH 4.0 acetate buffer at 37 °C. Reactions were monitored after the LLPS of HRP**.**

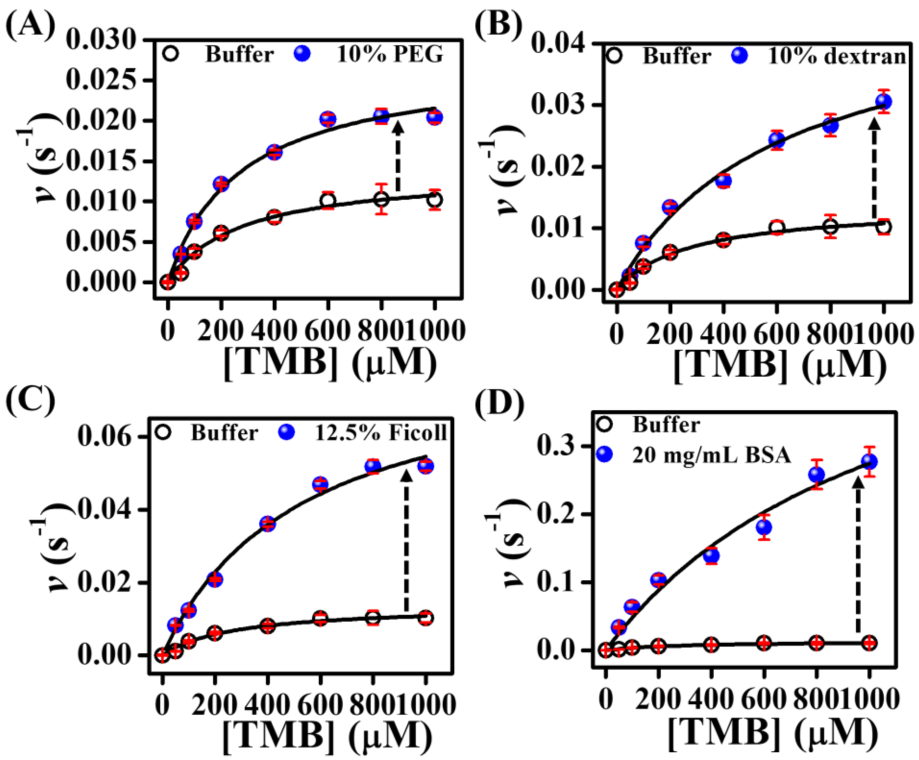


**Figure S19.** Michaelis-Menten plots of HRP as a function of TMB concentrations after the LLPS in pH 4.0 acetate buffer at 37 °C.

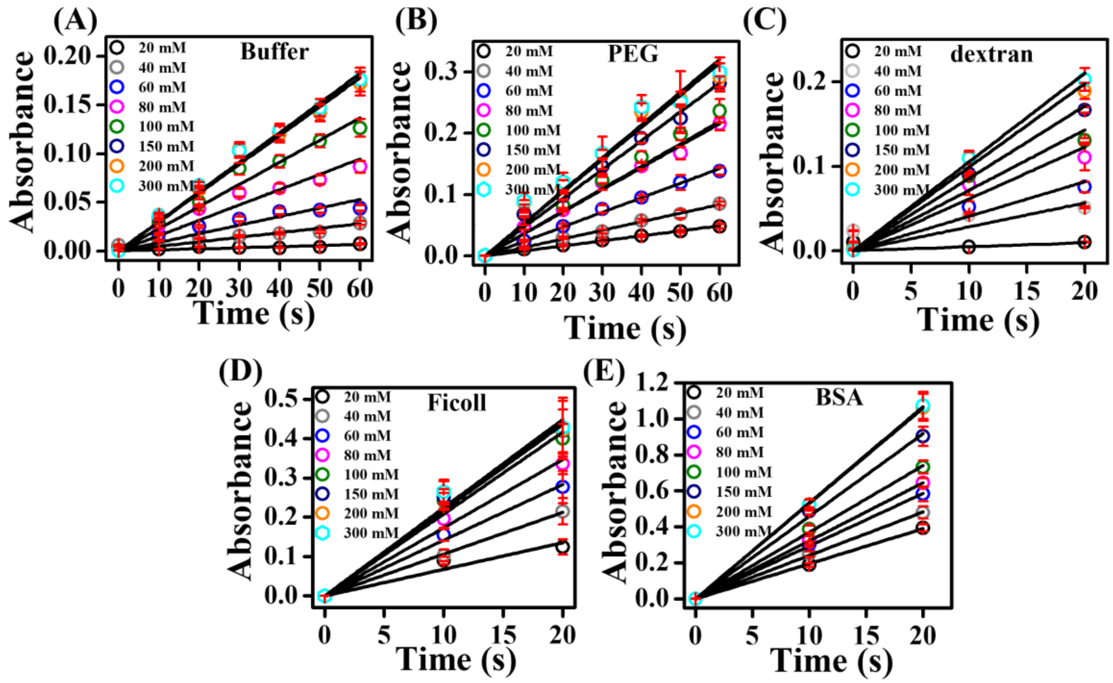


**Figure S20.** Plots of absorbance ($\lambda$ = 505 nm) versus time as a function of glucose concentrations in the absence and presence of different crowders in pH 7.4 PBS at 37 °C. Reactions were monitored after the LLPS of GOx**.**

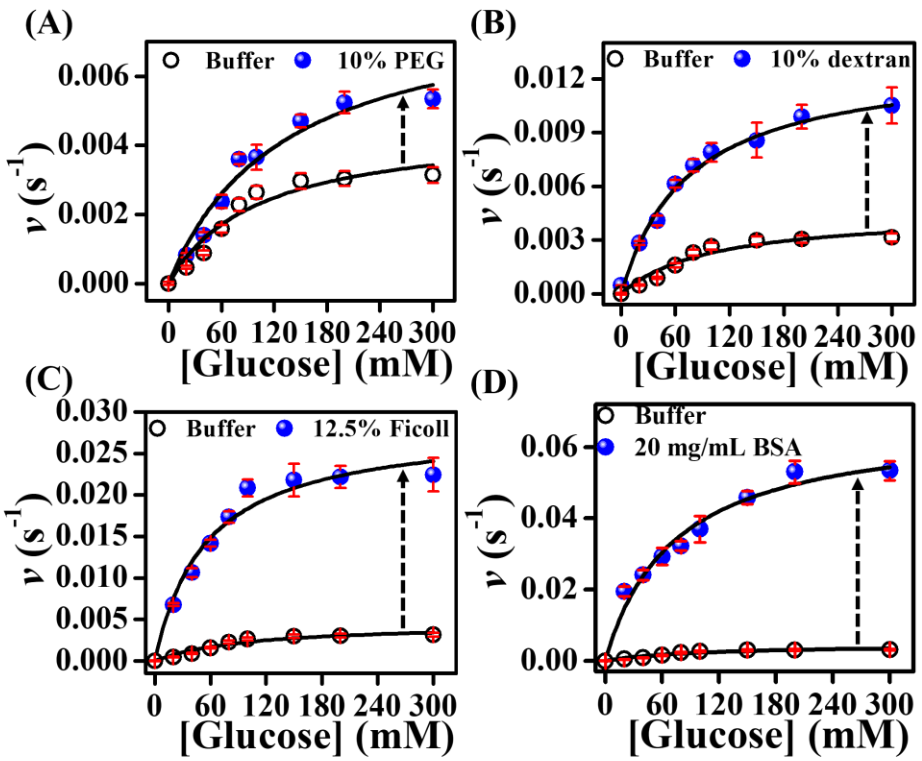


**Figure S21.** Michaelis-Menten plots after the LLPS of GOx in pH 7.4 PBS at 37 °C.

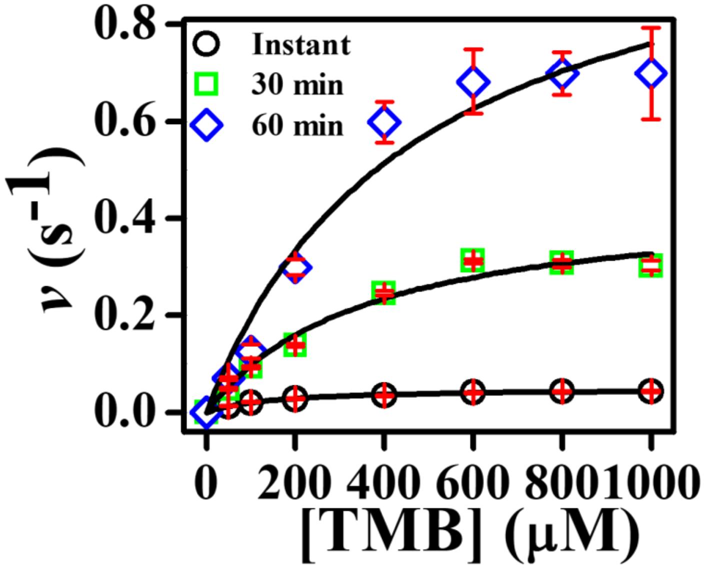


**Figure S22.** Michaelis-Menten plots after the LLPS of HRP in the presence of 10% dextran at different time intervals in pH 4.0 acetate buffer at 37 °C.

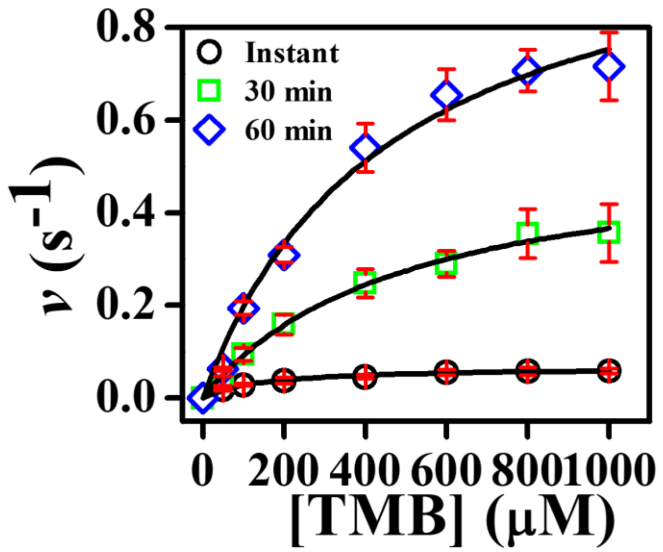


**Figure S23.** Michaelis-Menten plots after the LLPS of HRP in the presence of 12.5% Ficoll at different time intervals in pH 4.0 acetate buffer at 37 ℃.

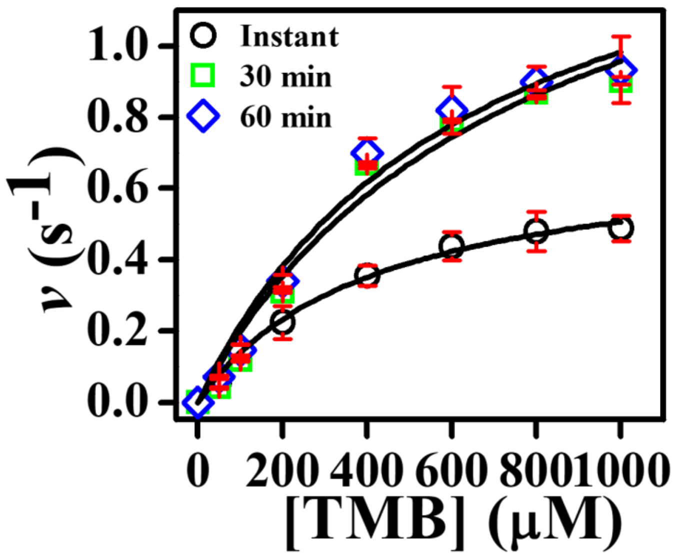


**Figure S24.** Michaelis-Menten plots after the LLPS of HRP in the presence of 20 mg/mL BSA at different time intervals in pH 4.0 acetate buffer at 37 °C.

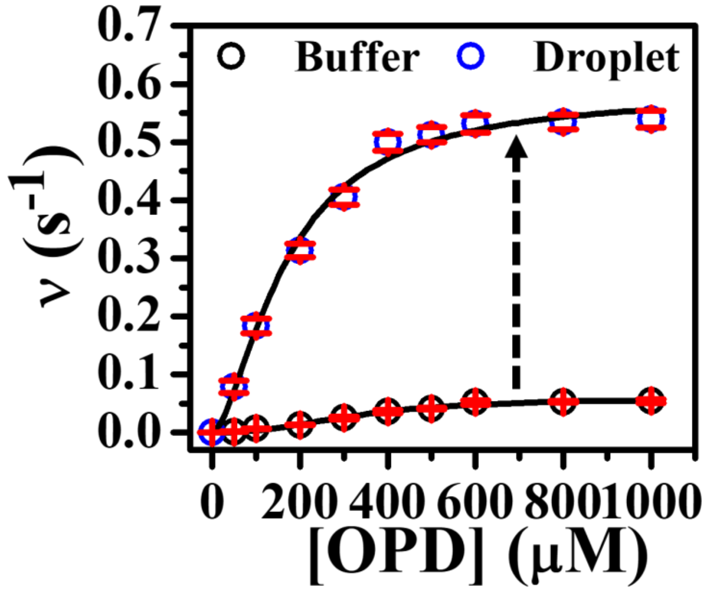


**Figure S25.** Michaelis-Menten plots of HRP as a function of OPD concentrations in the absence and presence of 10% PEG at 37 °C in pH 4.0 acetate buffer.

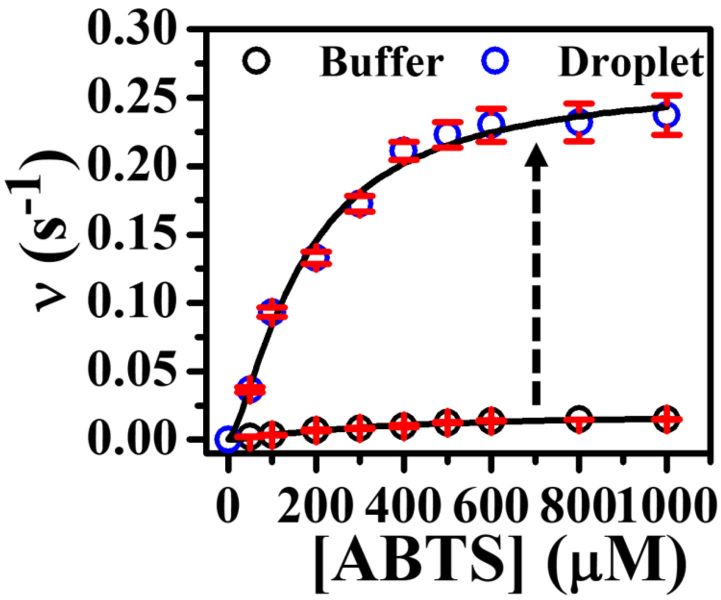


**Figure S26.** Michaelis-Menten plots of HRP as a function of ABTS concentrations in the absence and presence of 10% PEG at 37 °C in pH 4.0 acetate buffer.

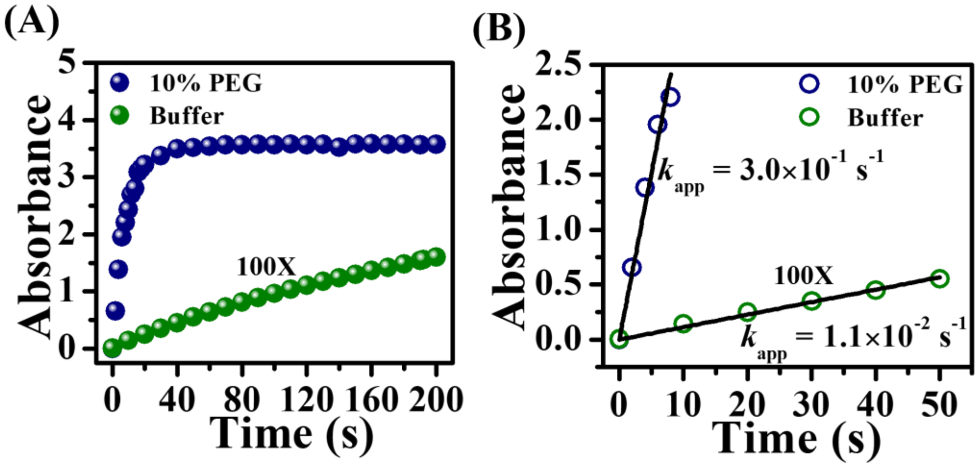


**Figure S27.** Changes in the (A) absorbance (λ = 650 nm) of ox-TMB and (B) $k_{app}$ of HRP in buffer with100-fold molar excess of enzymes and substrates along with those obtained in the presence of 10% PEG with 1X concentrations.

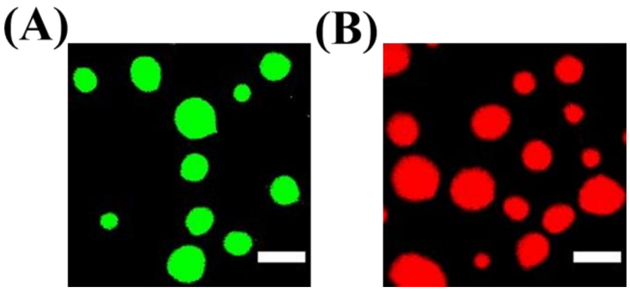


**Figure S28.** CLSM images of (A) FITC-labeled trypsin, and (B) RBITC-labeled alcohol dehydrogenase in the presence of 10% PEG in pH 7.4 PBS. The scale bars correspond to 5 μm.
